# PhyloForge: Unifying micro and macro evolution with comprehensive genomic signals

**DOI:** 10.1101/2024.03.06.583656

**Authors:** Ya Wang, Wei Dong, Yufan Liang, Weiwei Lin, Fei Chen

**Affiliations:** National Key Laboratory for Tropical Crop Breeding, Hainan University, College of breeding and multiplication, Sanya Institute of Breeding and Multiplication, Sanya 572025, China; College of tropical agriculture and forestry, Hainan University; Hospital of Stomatology, Guanghua School of Stomatology, Guangdong Provincial Key Laboratory of Stomatology, Sun Yat - Sen University, Guangzhou 510055, China

**Keywords:** PhyloForge, microevolution, macroevolution, phylogenomic signals

## Abstract

With the explosive growth of biological data, the dimensions of phylogenetic research have expanded to encompass various aspects, including the study of large-scale populations at the microevolutionary level and comparisons between different species or taxonomic units at the macroevolutionary level. Traditional phylogenetic tools often struggle to handle the diverse and complex data required for these different evolutionary scales. In response to this challenge, we introduce PhyloForge-a robust tool designed to seamlessly integrate the demands of both micro- and macro-evolution, comprehensively utilizing diverse phylogenomic signals, such as genes, SNPs, structural variations, as well as mitochondrial and chloroplast genomes. PhyloForge’s groundbreaking innovation lies in its capability to seamlessly integrate multiple phylogenomic signals, enabling unified analysis of multidimensional genomic data. This unique feature empowers researchers to gain a more comprehensive understanding of diverse aspects of biological evolution. PhyloForge not only provides highly customizable analysis tools for experienced researchers but also features an intuitively designed interface, facilitating effortless phylogenetic analysis for beginners. Extensive testing across various domains, including animals, plants, and fungi, attests to its broad applicability in the field of phylogenetics. In summary, the developmental background and innovative features of PhyloForge position it with significant potential in the era of large-scale genomics, offering a new perspective and toolset for a deeper understanding of the evolution of life.

## 1, Introductions

Phylogenetics, the cornerstone of evolutionary research, has progressively evolved into an interdisciplinary field of significant theoretical and practical implications. This domain encompasses data collection and analysis, software development, and phenotype association analysis. For instance, population genetics leverages extensive SNP data to examine disease genetic patterns. The phylogenetic status of various plants, including magnolia^1^, grape^2^, and waterlily^3^, is investigated using numerous low-copy genes. The origin and rapid evolution of the novel coronavirus are studied using whole genome sequences. In recent years, the swift advancement of DNA sequencing technology has led to an accumulation of substantial data for phylogenetic research^4^. Scientists have extracted a series of different types of phylogenetic signals from the genome, such as SNPs, low-copy genes, and small genomes like chloroplasts, mitochondrial genomes^5^, and viral genomes. To delineate the phylogenetic relationships of different taxa, it is essential to integrate vast amounts of data and various types of phylogenetic data due to the differences in evolutionary scales. This integration facilitates the completion of complex phylogenetic analysis.

There are some macroscopic or microscopic phylogenetic analysis tools, such as MEGA^6^, PAML^7^, phylobayes^8^. In recent years, the field of bioinformatics has seen a surge in the development of sophisticated software tools designed to address the complex challenges researchers face when analyzing biological data. These tools, which include ImageGP, Majorbio Cloud^9^, TCM-Suite^10^, and TBtools^11^, have revolutionized the way researchers approach data analysis. These software tools are specifically designed to streamline the process of biological data analysis, allowing researchers to delve deeper into their areas of interest without the need to become proficient in programming. This shift in focus from data management to data interpretation has significantly enhanced the efficiency and effectiveness of biological research. Moreover, these tools have democratized the field of bioinformatics, making it more accessible to researchers from diverse backgrounds. By simplifying the process of data analysis, these tools have enabled researchers to concentrate on formulating and testing hypotheses, thereby accelerating the pace of discovery in numerous fields of biological research. In essence, the advent of these bioinformatics analysis software tools has marked a significant milestone in the field of biological research, empowering researchers to transcend the traditional boundaries of data analysis and explore new frontiers in their respective fields.

Phylogenetic analysis still lacks tools that integrate macroscopic and microscopic aspects. The fundamental operating principle of phylogenetic analysis tools is to elucidate the phylogenetic relationships among sequences. This is achieved by aligning biological sequences, computing evolutionary distances, constructing phylogenetic trees, and subsequently evaluating and optimizing these trees. Currently, the field is equipped with specialized tools such as Phylosuite^12^ and MEGA^6^, which are tailored for specific tasks. However, a comprehensive tool that can analyze multiple types of phylogenetic signals is yet to be developed. In essence, there is a lack of a unified tool that can simultaneously analyze microevolution and macroevolution. Given the varying scales of research, the data analyzed also differs. Therefore, there is a pressing need for a comprehensive tool that can integrate various data types for analysis, thereby facilitating a more efficient and effective research process.

This study is designed to bridge the existing gap in tools for macroevolution and microevolution, with an emphasis on expediting the process of tree drawing and simplifying intricate procedures. We have pioneered the development of a comprehensive tool, PhyloForge. Our objective is to establish a unified platform that amalgamates research in macroevolution, microevolution, and sequence evolution, thereby facilitating swift analysis of extensive data and diverse data types.

## 2, Overview

The PhyloForge tool consists of three major components:

1. Single Nucleotide Polymorphisms (SNPs)^13^ and Structural Variations (SVs)^14^ are two sequence features that exhibit a whole-genome distribution. By employing two distinct methodologies, they elucidate systematic relationships and are utilized for microevolutionary studies at the subspecies level.
2. Three methods are used to measure phylogenetic relationships through low-copy nuclear genes and mitochondrial or chloroplast genomes^15,16^, focusing on macroevolution above the species level.
3. Beyond the study of phylogenetics at the cellular, individual, or population level, it is imperative to investigate systematic relationships at the sequence level. To cater to this requirement, we have incorporated the most advanced and superior quality tools for sequence alignment and phylogenetic tree construction. This approach ensures a comprehensive analysis at the sequence level.

The aforementioned tools have been seamlessly integrated into the PhyloForge toolkit. Given the substantial computational demands associated with calculating macroevolution and microevolution, these tasks surpass the computational capabilities of personal computers or small servers. Consequently, PhyloForge has been designed as a command-line tool, with Linux serving as its foundational operating system. To facilitate users in swiftly mastering the input and output files along with the associated parameters, we provide comprehensive documentation and examples. Additionally, we offer extensive video and text materials for related tutorials. All these parameters can be accessed through the ‘—help’ command. Comprehensive documentation and case information are readily available on GitHub.

## 3. Program pipeline

At the heart of PhyloForge software lies the utilization of various phylogenetic signals at the whole genome level to construct phylogenetic relationships. This pipeline encompasses a series of Python modules, incorporating both external software and code that we have financed for development.

The entire process primarily comprises five stages: (1) File preparation, (2) Data processing, (3) Sequence alignment and correction, (4) Evolutionary tree establishment, (5) Output of files in various formats.

As a command-line tool, PhyloForge primarily employs ten primary parameters (-l, -m, -o, -w, -s, -S, -g, -grc, -gsc, -h) and a multitude of secondary parameters. This ensures that each step of the various phylogenetic analyses is executed with precision and rigor (Table 1).

**Table 1.**
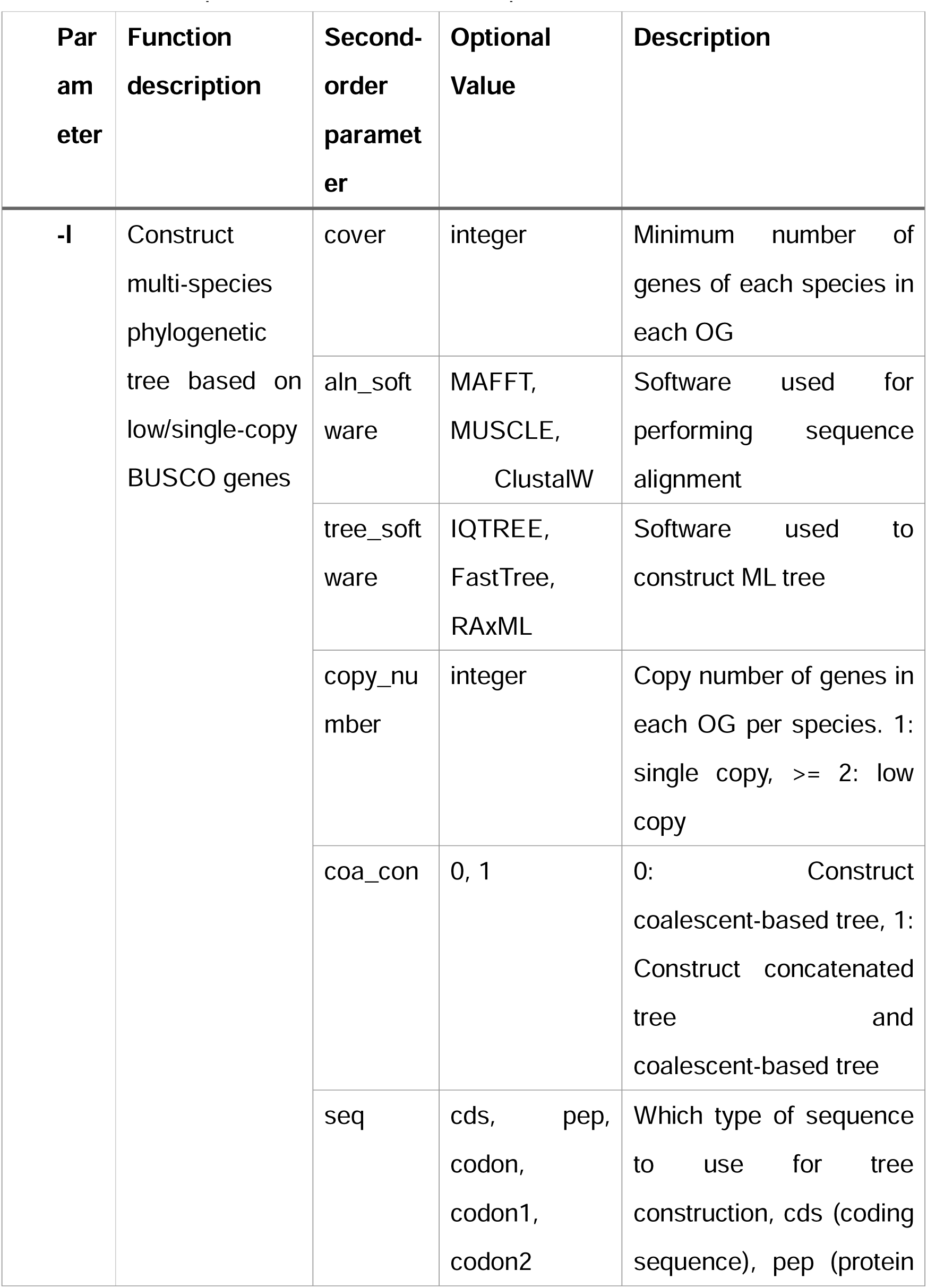
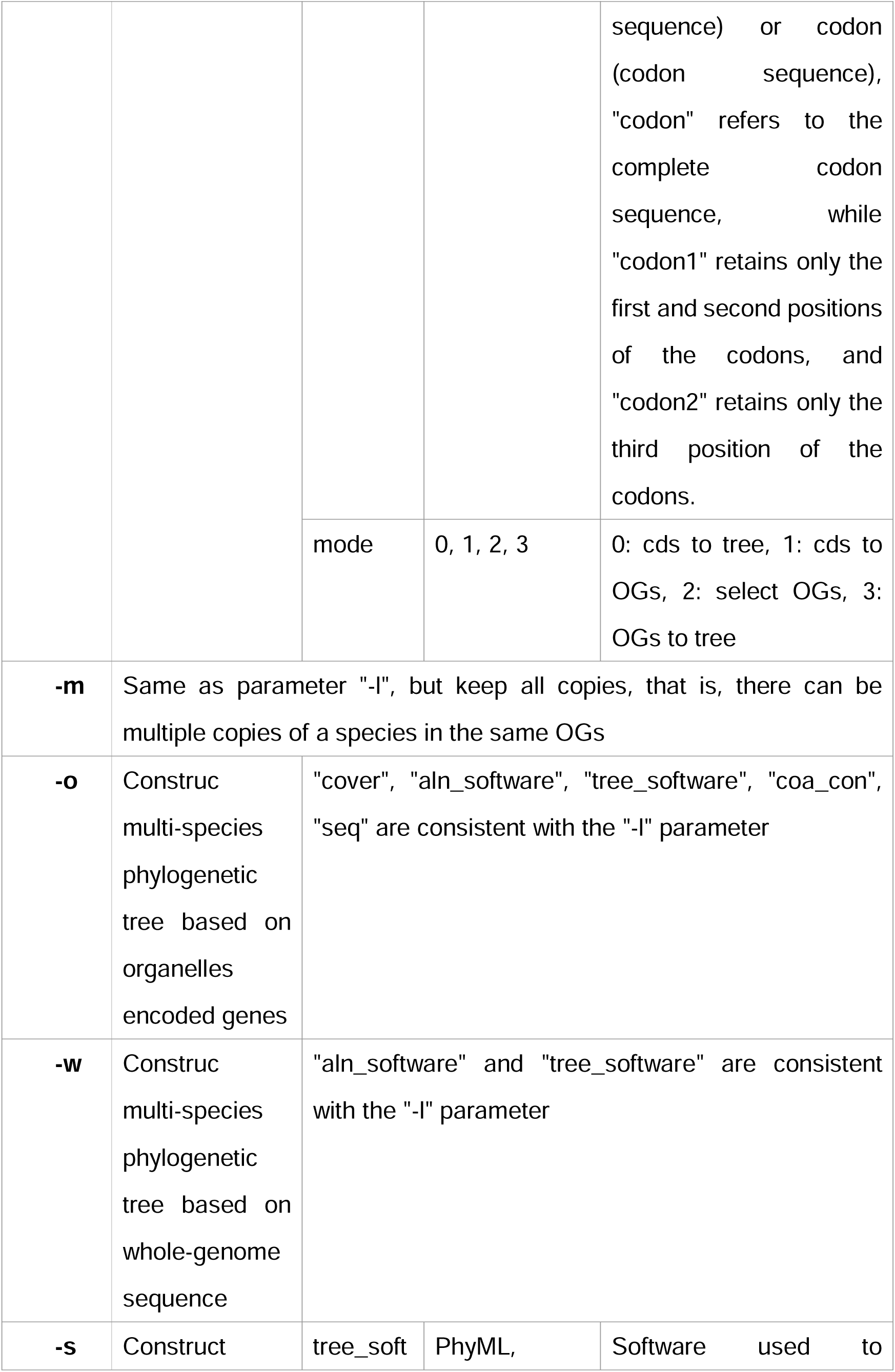
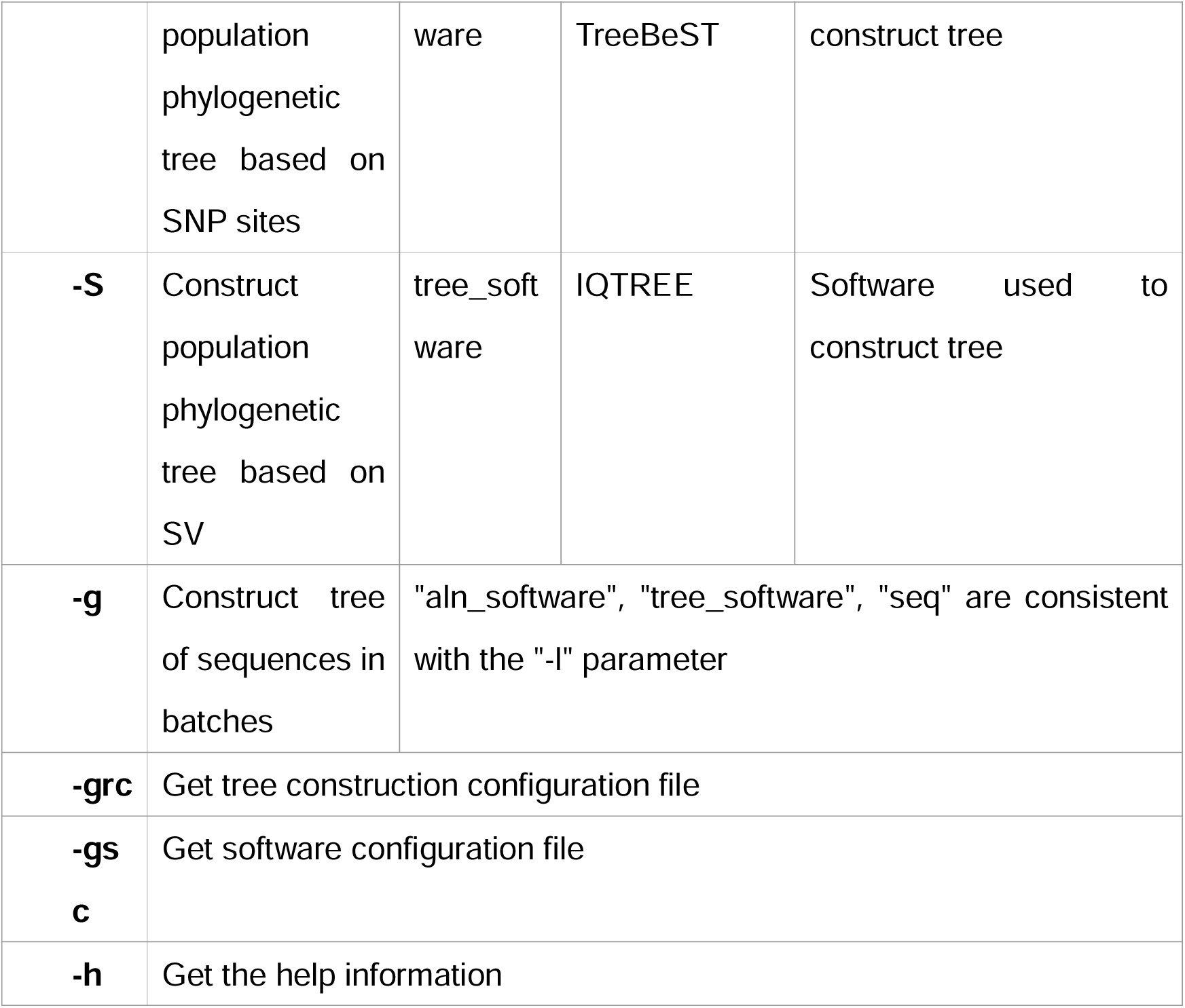
The parameters and their descriptions of TreeTool.

### Microevolution level

Single Nucleotide Polymorphisms (SNPs) are densely distributed phylogenetic signals at the whole genome level that have been extensively studied and widely utilized. They are apt for inferring the degree of genetic differentiation among populations, identifying the extent of gene flow, and can also be employed to estimate the time of differentiation among different populations or species^17–19^. Owing to the substantial computational resources and server support required for SNP calling, PhyloForge employs input .vcf files, which comprise a collection of filtered, high-quality SNPs, to directly conduct phylogenetic analysis. Initially, we convert the genotypes of different individuals into nucleic acid sequences. Subsequently, we utilize efficient and swift tree construction software, namely TreeBest^20^ and PhyML^21^, to infer the phylogenetic tree.

In the realm of population genetic research, both Single Nucleotide Polymorphisms (SNPs) ^13,18^ and Structural Variations (SVs) ^17,22^ are taken into account by researchers to gain a comprehensive understanding of genetic variation and the evolutionary history of the genome. Both these elements play a pivotal role in unveiling biological diversity and evolutionary processes at various levels. As a crucial supplement to SNP-based phylogenetics, we have pioneered the development of a phylogenetic research process centered on SVs^23–27^. This process involves the extraction of the most significant and easily distinguishable insertions and deletions. These are then converted into a 0-1 distribution, which is subsequently transformed into a complete 0-1 matrix. Finally, IQTREE^28^ is employed to infer the phylogenetic tree.

### Macroevolution level

For phylogenetic analysis conducted above the species level, the substantial evolutionary differences necessitate the use of relatively conservative low-copy nuclear genes to illustrate the evolutionary relationships between species. This method is widely recognized and is applicable to a broad scale, encompassing the entire terrestrial plant group with a history spanning over 400 million years, as well as all types of fungi. PhyloForge supports users in inputting a series of plant CDS sequences. Following the extraction of all high-quality low-copy nuclear genes, users have the autonomy to select the species coverage and copy number in each Orthologous Group (OG). Through sequence alignment and trimming, users can opt to use multiple tree-building software to infer phylogenetic relationships. To facilitate a more detailed sequence analysis, users have the flexibility to choose the final sequence used to infer phylogenetics. Users can select the original CDS sequence, the protein sequence translated from CDS, or the codon sequence converted from CDS and protein. For codon sequences, users can choose the three-position codon of 1st+2nd+3rd positions, or they can select the codon of 1st+2nd positions or only the 3rd position (Table 1). This versatility aids in deciphering complex evolutionary relationships.

In the context of macroevolution, plants often encounter numerous conflicts, such as nuclear conflicts instigated by gene introgression and incomplete lineage sorting. Consequently, phylogenetics based on organelle genomes, in addition to those based on nuclear genes, play a pivotal role in unraveling complex evolutionary events in the evolutionary history of species^15,16^. In light of this, we have developed two distinct modules within PhyloForge for phylogenetic analysis predicated on organelle genome data. The first module, referred to as the ‘Complete Genome Sequence Analysis’, infers phylogenetic relationships based on the complete genome sequence of organelles. The second module, termed the ‘Organelle-Encoded Gene Analysis’, infers phylogenetic relationships based on organelle-encoded genes. Users can select the appropriate module based on the biological problems they aim to solve and the degree of kinship between species.

### Sequence level

Sequence comparison is a standard method and a crucial foundation for studying gene function. Given the necessity for an exceedingly precise and accurate comparison between sequences, PhyloForge has the capability to utilize multiple tools such as MAFFT^29^, MUSCLE^30^, and ClustalW^31^ for pairwise sequence alignment and multiple sequence alignment. Concurrently, it can opt to use IQTREE^28^, FastTree^32^, and RAxML^33^ for the construction of evolutionary trees. This versatility aids in advancing phylogenetic analysis of viruses and plasmids, as well as sequences that extend beyond the individual level.

### Quality Control

Currently, the paramount factor influencing the precision of phylogenetics is data quality control. PhyloForge executes a variety of quality controls tailored for different types of signals. For instance, users have the flexibility to set parameters such as gene copy number and coverage to fulfill specific requirements. In the case of sequence alignment, trimming operations are necessitated. Moreover, users have the autonomy to independently select sequence alignment software, tree-building software, and sequences for tree construction, among other options. Users can freely conduct quality control in accordance with their individual needs to cater to diverse analysis requirements.

### Output

PhyloForge primarily generates tree files, encompassing ‘.nwk’ and ‘.nhx’ files. Additionally, it has the capability to output crucial intermediate files. These include matrices derived from Structural Variation (SV) data, fasta sequences converted from Single Nucleotide Polymorphism (SNP) sites, and orthogroup lists, among others. These resources are invaluable for professional users, facilitating continuous improvement and correction in their work.

## 4, Applications

### 4.1 Systematic classification of higher plants

We have meticulously selected the genomes of angiosperms from a diverse range of taxa to examine the large-scale phylogeny of higher plants. For such macroevolution spanning a scale of 200 million years, our strategic approach involves the utilization of all Coding Sequence (CDS) files to mine low-copy nuclear genes as phylogenetic signals^34–37^. We have chosen 76 flowering plants and screened a total of 230 Orthologous Groups (OGs) (Fig 3, Table S1). We employed the BUSCO method, a technique not widely used hitherto, to identify Low-Copy Nuclear genes (LCNs). Consequently, we performed Gene Ontology (GO) and Kyoto Encyclopedia of Genes and Genomes (KEGG) annotation analysis on these 230 low copy genes identified in Amborella trichopoda. Our results revealed that out of the 230 genes, 127 were annotated with GO terms, while 154 genes were annotated with KEGG pathways (Fig S1). These genes were predominantly involved in critical biological processes such as nitrogen compound metabolic process and DNA metabolic process (Biological Process). This substantiates that the stringent parameters of PhyloForge can effectively screen suitable phylogenetic signals at the whole genome level, and are applicable to the macroscopic evolution of plants on a scale of 200 million years. The inclusion of the results of phylogenetic development should be considered.

### 4.2 Systematic classification of higher animals

In contrast to plants, animals, with the exception of a few fish species^38,39^, have generally not undergone polyploidization events since the Cambrian period. While PhyloForge is applicable to plants that have experienced multiple polyploidizations^40–42^, we have also explored its application in animal phylogenetics. We selected 18 primates with sequenced genomes for this purpose (Table S2). The evolutionary tree output by PhyloForge exhibits robust support, with a posterior probability (PP) of 1 or a bootstrapping score (BS) of 100 for each branch. Furthermore, the topology of the tree aligns perfectly with widely accepted reports, demonstrating its effective application in the study of animal evolution.

### 4.3 Classification of fungi

Fungi represent the most diverse group among all eukaryotes, encompassing approximately 5 million species. To illustrate the capabilities of PhyloForge, we have utilized the phylogeny of Arbuscular Mycorrhizal (AM) fungi as an exemplar. Employing the genomes of 26 distinct AM fungi (Table S3), PhyloForge can yield robust phylogenetic results (Table S4), this includes the generation of clear and unambiguous branches, with each node receiving substantial support, denoted by a posterior probability (PP) of 1 or a bootstrapping score (BS) of 100. These results underscore the efficacy of PhyloForge as a valuable tool for conducting phylogenetic research within the realm of fungal studies.

### 4.4 Phylogeny of the Theaceae family based on chloroplast genomes

In our research, we utilized chloroplast genomes to investigate the evolution of the primary plants within the Theaceae family. Our analysis incorporated chloroplast genome data from 14 species of Theaceae and one species of Pyrolaceae. Our examination of the plastid genomes of these 14 Theaceae species and one Pyrolaceae species revealed that the phylogenetic tree, constructed based on organellar-encoded genes and whole-genome sequences, exhibited consistent topologies (as depicted in Fig 2D). These findings were generally in alignment with previous studies^43^. These results underscore that PhyloForge, in addition to handling phylogenetic analysis based on low-copy nuclear genes, also excels in conducting phylogenetic analysis predicated on organelles.

### 4.5 Phylogeny of tea plant populations based on SNPs

For taxonomic units at the subspecies level and below, Single Nucleotide Polymorphisms (SNPs) that are distributed at the genome level are commonly employed as phylogenetic signals. In our study, we utilized SNP data from 60 tea tree populations for testing purposes. The phylogenetic analysis of these 60 tea plants, based on SNP data, suggested that these plants could be segregated into two primary evolutionary branches (as depicted in Fig 2E). This finding was in alignment with the results of the population structure analysis (Fig S3). These results underscore that PhyloForge exhibits commendable accuracy in conducting SNP-based phylogenetic analysis.

### 4.6 SV signal-based population phylogeny

Structural variations, characterized by insertions, deletions, inversions, and translocations, serve as crucial observational indicators when comparing populations within species. They have emerged as a novel research field in recent years, largely due to advancements in long-read sequencing technology. In response to this, we have pioneered a new phylogenetic method predicated on Structural Variations (SVs), employing insertions and deletions as 1 signal and 0 signal respectively. We utilized data from 507 maize individuals to explore the application of these two structural variations^44^. At the whole genome level of maize, these two SVs exhibit high coverage on chromosomes. We transform these SVs into 0-1 phylogenetic signals, convert them into a 0-1 matrix, and ultimately translate them into a phylogenetic tree file. Upon examination, we observe that the maize phylogenetic tree exhibits a high degree of distinction and doesn’t lack ambiguous nodes. Additionally, we evaluated the discriminatory power of Phyloforge for Camellia sinensis using SV data from 136 individuals^45^. Our results demonstrated that Phyloforge exhibited a significant discrimination effect for Camellia sinensis. Consequently, it serves as a potent supplement to the evolutionary tree with Single Nucleotide Polymorphisms (SNPs) as the signal.

In comparison to analogous tools, PhyloForge significantly streamlines the intricate process of phylogenetic analysis. It offers a user-friendly interface that is faster and more efficient, thereby reducing the time required for analysis. Moreover, PhyloForge provides a comprehensive set of parameters, offering users a high degree of flexibility and control over their analyses. One of the standout features of PhyloForge is the inclusion of the newly developed innovative tool, Structural Variation (SV). This tool represents a significant advancement in the field of phylogenetic analysis, offering new possibilities for researchers. PhyloForge is designed to cater to a wide range of needs, from sequence analysis to macro analysis spanning over 500 million years of history, and micro analysis of population genetics. Whether you’re investigating the evolutionary history of a species or studying the genetic variation within a population, PhyloForge provides the tools and resources you need to conduct comprehensive and rigorous analyses (Table 2).

**Table 2.**
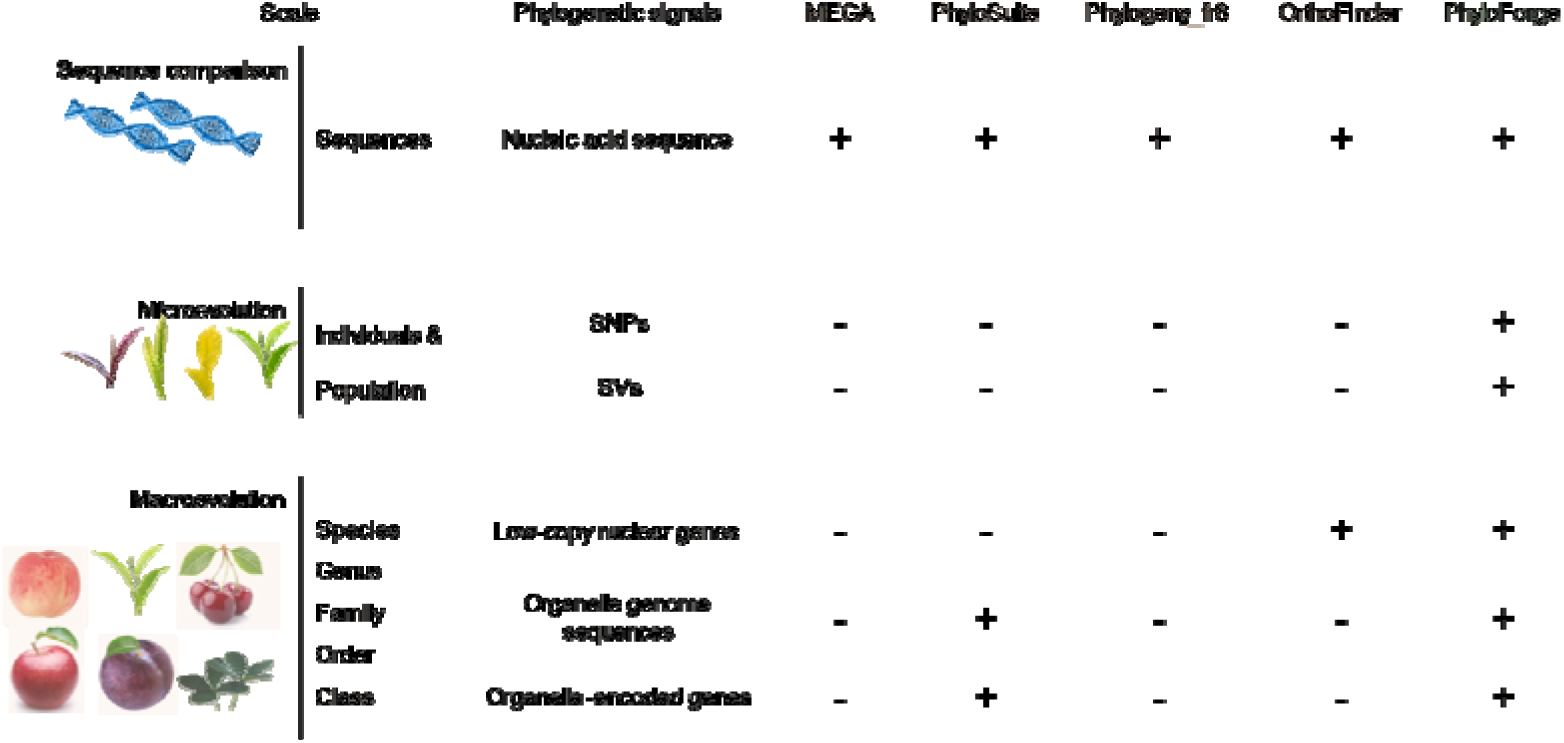
Comparison of PhyloForge with popular tools offering similar functionalities.

## 5, Discussion

The science of phylogenetics is dedicated to reconstructing the evolutionary relationships among organisms, and it serves as the foundation of biological research^46–49^. Accurate phylogenetic relationships are crucial prerequisites for analyzing the evolutionary trajectories of genes, genomes, and species^50–52^. Therefore, obtaining accurate phylogenetic relationships has always been a goal for biologists. From the past practice of classifying based on biological morphology, to the subsequent use of individual or a few organelle-encoded genes for phylogenetic analysis, to the current explosion of sequencing data, it is now mainstream in phylogenetic research to obtain more accurate phylogenetic relationships based on whole genome sequences and to analyze complex evolutionary events that affect the stability of phylogenetic relationships.

Analyzing the evolution of simple sequences, resolving population-level evolutionary relationships based on whole-genome SNPs or SVs, and deciphering cytonuclear conflicts based on organelle genome sequences and nuclear-encoded genes are among the most common and fundamental analyses in phylogenetic research. These are also key foundations for understanding biological evolution. However, the complex analysis process poses a significant challenge to researchers in the field of biology. For instance, current methods for phylogenetic analysis based on orthologous single-copy nuclear genes often involve a series of complex processes, including the selection of suitable orthologous groups (OGs), sequence alignment, trimming, and tree construction. In addition to a solid foundation in biology, a certain level of computer literacy is also required. Furthermore, there are often many uncertainties in this process, such as the inability to select suitable OGs.

Given the importance of phylogenetic research and the challenges currently faced in phylogenetic analysis, we have developed a comprehensive phylogenetic toolkit called PhyloForge. This toolkit is fully functional, integrating common methods of phylogenetic analysis to meet different levels of phylogenetic analysis needs. It can perform all analyses of sequence evolution, including microevolution and macroevolution of species evolution, which no other tool has achieved so far. In comparison to similar phylogenetic tools in the past, we have further simplified the user experience with PhyloForge. Users can directly provide input files (.fa/.vcf) and obtain phylogenetic results (.nwk/.NHX/.treefile) in one step. This solves the problem of independent functionalities in tools like MEGA and PhyloSuite, eliminating the need for manual handling of any intermediate result files. This is particularly important for researchers who do not have a strong programming background or are unfamiliar with the phylogenetic analysis process. Reliability is another major advantage of PhyloForge. The tool is developed based on the most commonly used and highly recognized phylogenetic analysis methods. The data test results of each module demonstrate extremely high reliability.

The functionality, ease of use, and reliability of PhyloForge make it highly suitable for both novices and experienced users in phylogenetic analysis, aiding them in their phylogenetic research endeavors.

## 6, Conclusion

In summary, the test results collectively underscore the efficacy of our tool, PhyloForge. It is not only user-friendly, offering a high degree of flexibility in terms of parameter settings, but also exhibits exceptional accuracy in its operations. This amalgamation of user-centric design and precision makes PhyloForge a convenient and reliable choice for researchers engaged in phylogenetic analysis. Its robust performance ensures that it can handle a wide array of analytical tasks, thereby serving as a comprehensive solution for phylogenetic research.

## 7, Methods

The workflow depicted in Figure 1 illustrates the fundamental steps involved in this tool for constructing a phylogenetic tree. We have mainly designed three phylogenetic analysis modules.

**Figure 1.**
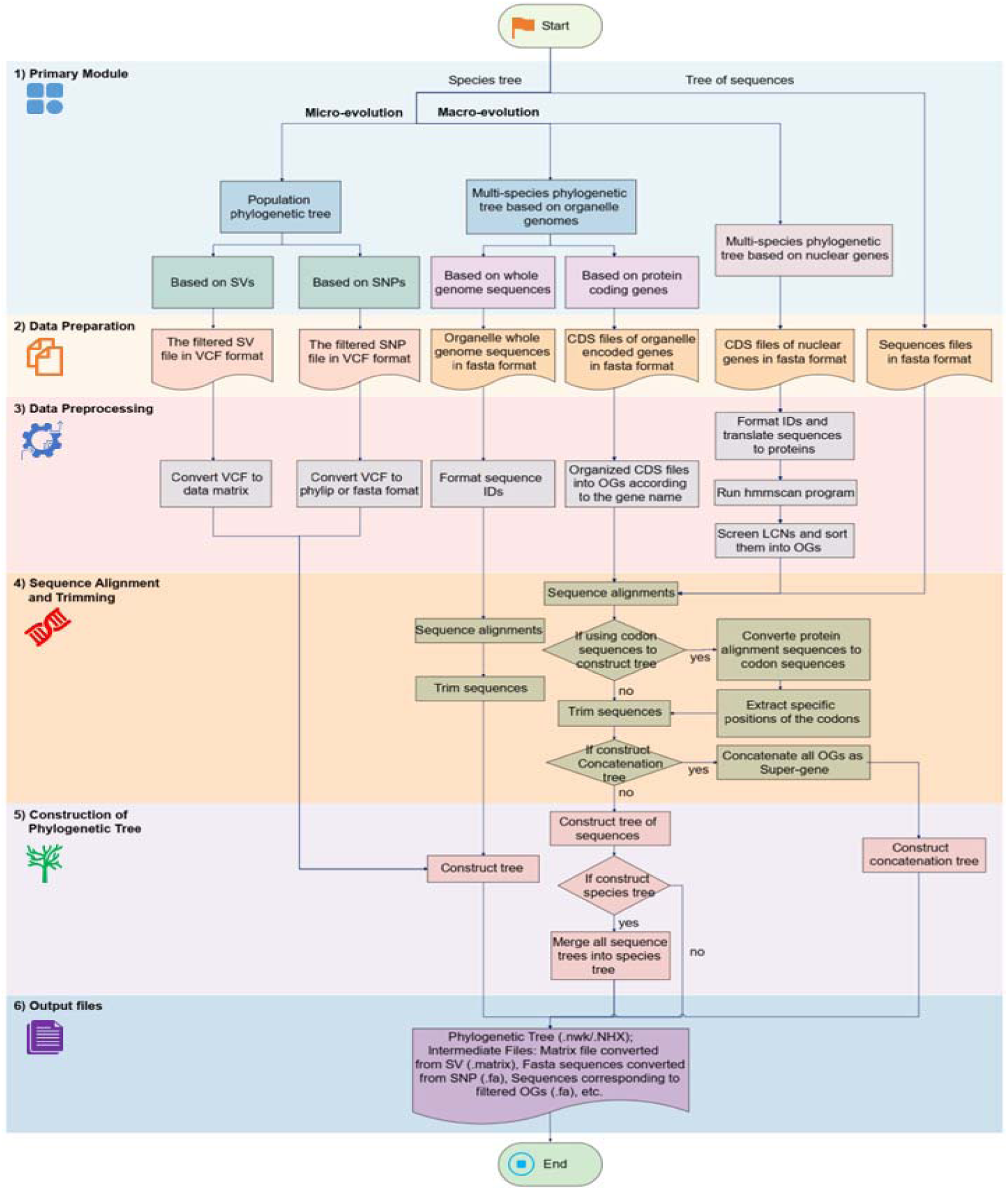
The pipeline of the software. PhyloForge consists of three main functions, phylogenetic inference at micro-evolution, macro-evolution, and the sequence level comparison. PhyloForge process these different kinds of phylogenetic signals and convert them into matrix, then construct and output the phylogenetic trees.

**Figure 2.**
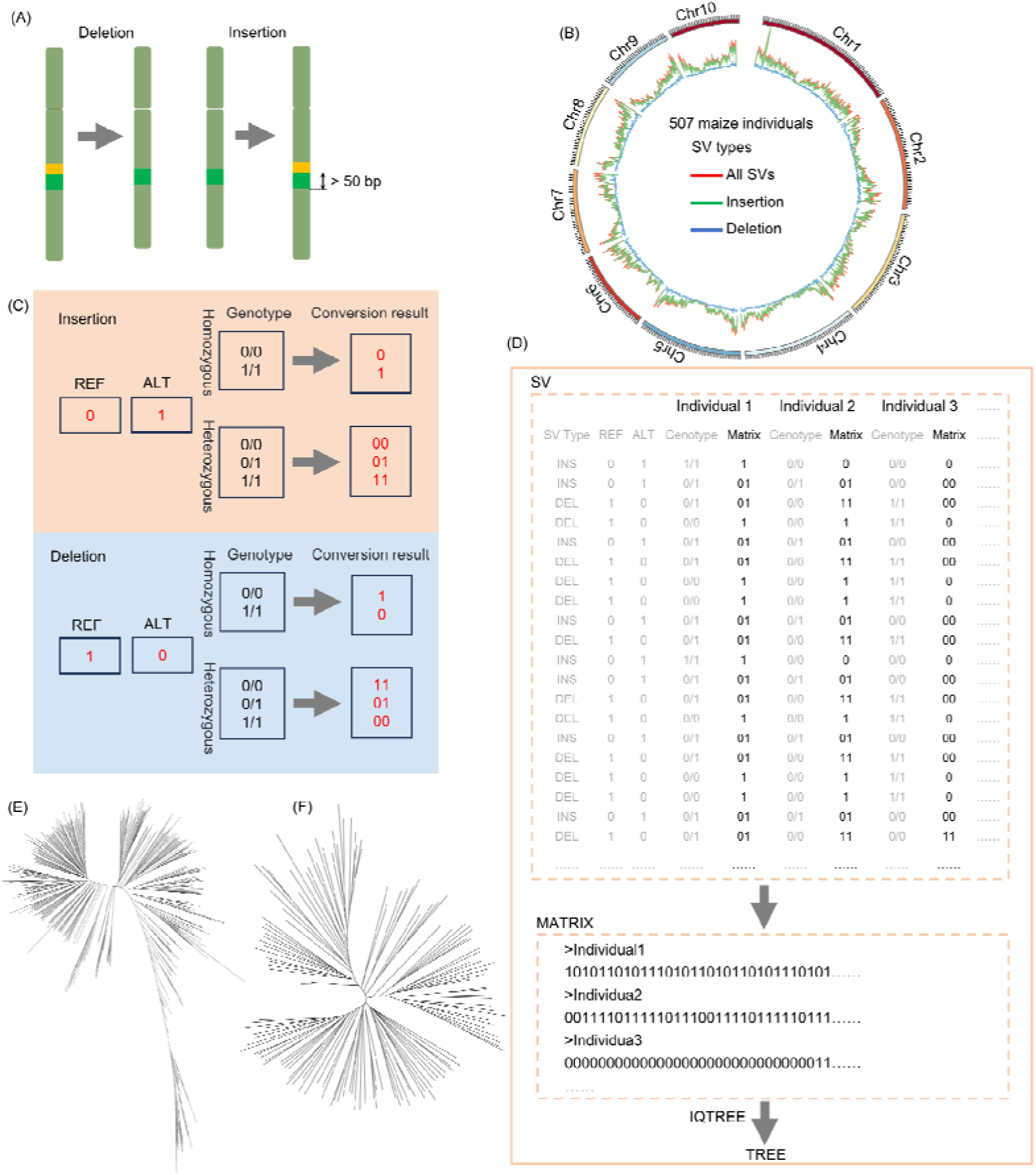
Principles of microphylogenetic analysis based on SV and its robust application. (A) Introduction to two types of structural variations (SVs): deletions (DELs) and insertions (INSs). (B) Abundant SVs dominated by DELs and INSs in the population, taking maize Chr1 as an example. (C) Principles for converting DELs and INSs into matrix. (D) Process of constructing a tree from SVs. (E) A phylogenetic tree of 507 maize individuals constructed based on the above principles using SVs. (F) A phylogenetic of 136 tea plants (*Camellia sinensis*) constructed based on the above principles using SVs.

**Figure 3.**
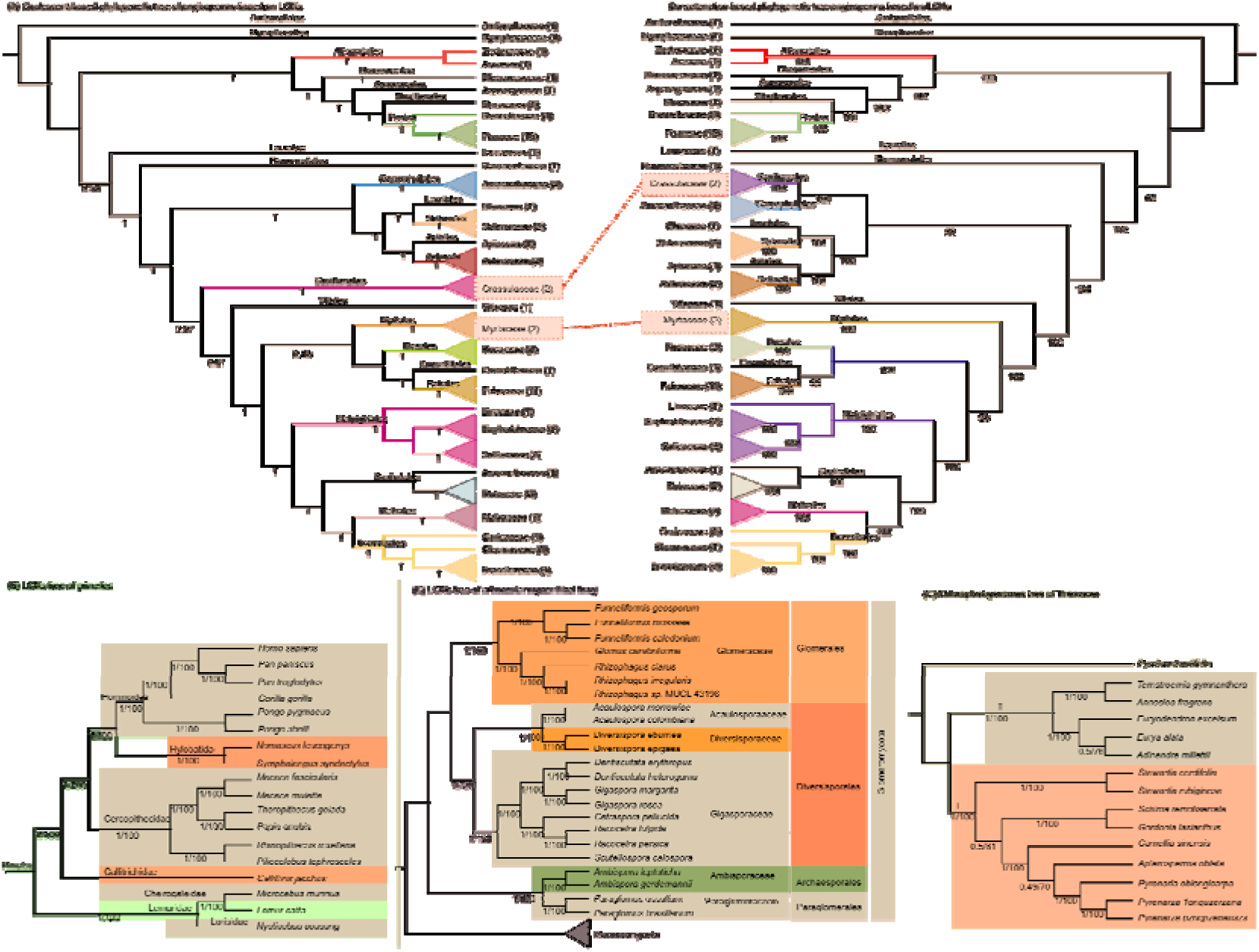
Results of macroscopic phylogenetic analysis utilizing PhyloForge. (A) Depicts a coalescent-based (left) and concatenation-based (right) phylogenetic tree, derived from BUSCO genes of 76 angiosperm species across 31 families. The number beneath the branch represents the local posterior probability (PP) for the coalescent-based tree, or the bootstrapping support (BS) for the concatenation-based tree. (B) Illustrates a phylogenetic tree based on BUSCO genes of 18 primate species within 7 families. The number beneath the branch represents PP/BS. (C) It shows a phylogenetic tree based on BUSCO genes of arbuscular mycorrhizal fungi. The number beneath the branch represents PP/BS. (D) It presents a phylogenetic tree of Theaceae based on Coding DNA Sequences (CDS) (Coalescent-based) and whole genome sequences of chloroplast genomes. The number beneath the branch represents PP/BS. This passage has been revised for a more formal tone.

### 7.1 macro-scale multi-species phylogenetic analysis

For macro-scale multi-species phylogenetic analysis, we have developed two modules. (1) Based on low-copy BUSCO (Benchmarking Universal Single-Copy Orthologs) genes for multi-species phylogenetic analysis. This module primarily involves identifying BUSCO genes in the species, then retaining low/single-copy BUSCO genes for phylogenetic analysis. The main workflow includes using HMMER hmmscan to identify species-specific BUSCO genes, selecting low/single-copy BUSCO genes to organize OGs (Orthologs groups), aligning and trimming sequences, and then constructing trees. (2) For multi-species phylogenetic analysis based on organelle genomes, we have designed two modes: based on whole organelle genome sequences and based on organelle-encoded gene CDS sequences. The construction of the phylogenetic tree based on organelle-encoded genes entails concatenating or parallelizing sequences, aligning and trimming them, and subsequently constructing the phylogenetic tree. Similarly, the construction of the phylogenetic tree based on whole genome sequences involves aligning and trimming the nucleotide sequences of the entire genome, followed by tree construction.

### 7.2 micro-scale population phylogenetics

For the analysis of micro-scale population phylogenetics, we have developed two modules. The first module is designed for phylogenetic analysis based on SNP sites. It involves the conversion of VCF format files into fasta or phylip format nucleotide sequences using IUPAC Ambiguity Codes, followed by the construction of phylogenetic trees based on these sequences. Additionally, we have designed a module for phylogenetic analysis based on structural variations (SVs). This module primarily converts insertions and deletions, the two types of structural variations, into matrices, which are then used to construct phylogenetic trees.

### 7.3 evolutionary trees at the sequence level

In addition, we have developed a multi-sequence phylogenetic analysis module, which can perform phylogenetic analysis of a large number of sequences in multiple processes and complete the construction of a large number of gene trees at one time.

To implement this workflow, we have utilized the Python programming language, and all the source code has been uploaded to GitHub(https://github.com/wangyayaya/PhyloForge).

## Conflict of interest

The authors declared none conflict of interest.

## Author contributions

F.C. designed and led this study. Y.W., W.D., Y.L., W.L. performed the whole programming and benchmark. Y.W., Y.L., W.L., and F.C. wrote the manuscript. All the authors approved the final manuscript.

## Acknowledgement

This work was supported by the National Natural Science Foundation of China (32172614), Hainan Province Science and Technology Special Fund (ZDYF2023XDNY050), Hainan Provincial Natural Science Foundation of China (324RC452), and the Project of National Key Laboratory for Tropical Crop Breeding (NO. NKLTCB202337).

## Supplementary materials

**Supplement figure 1.**
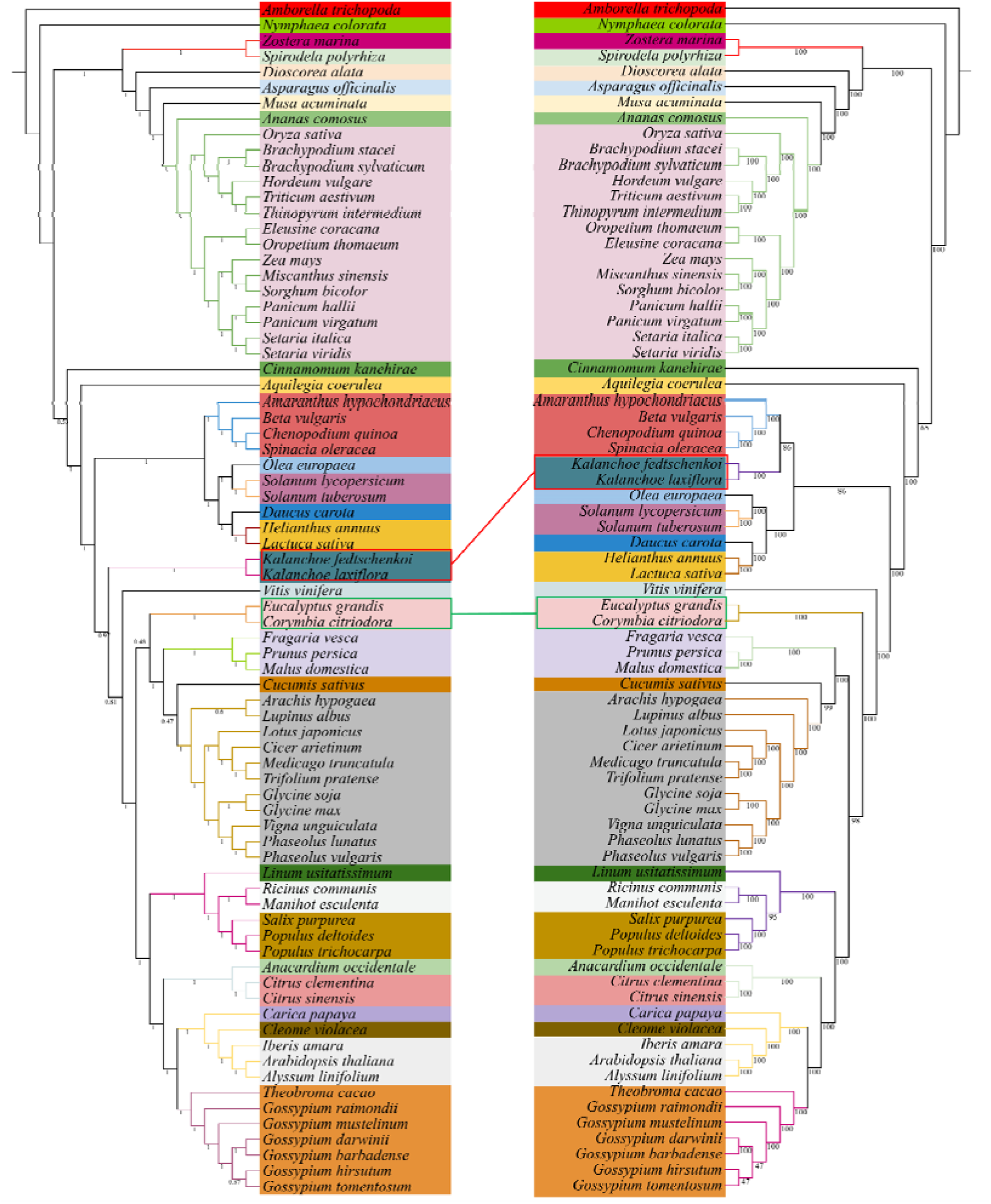
Coalescent-based (left) and concatenation-based (right) phylogenetic tree of 76 angiosperm species.

**Supplemental figure 2.**
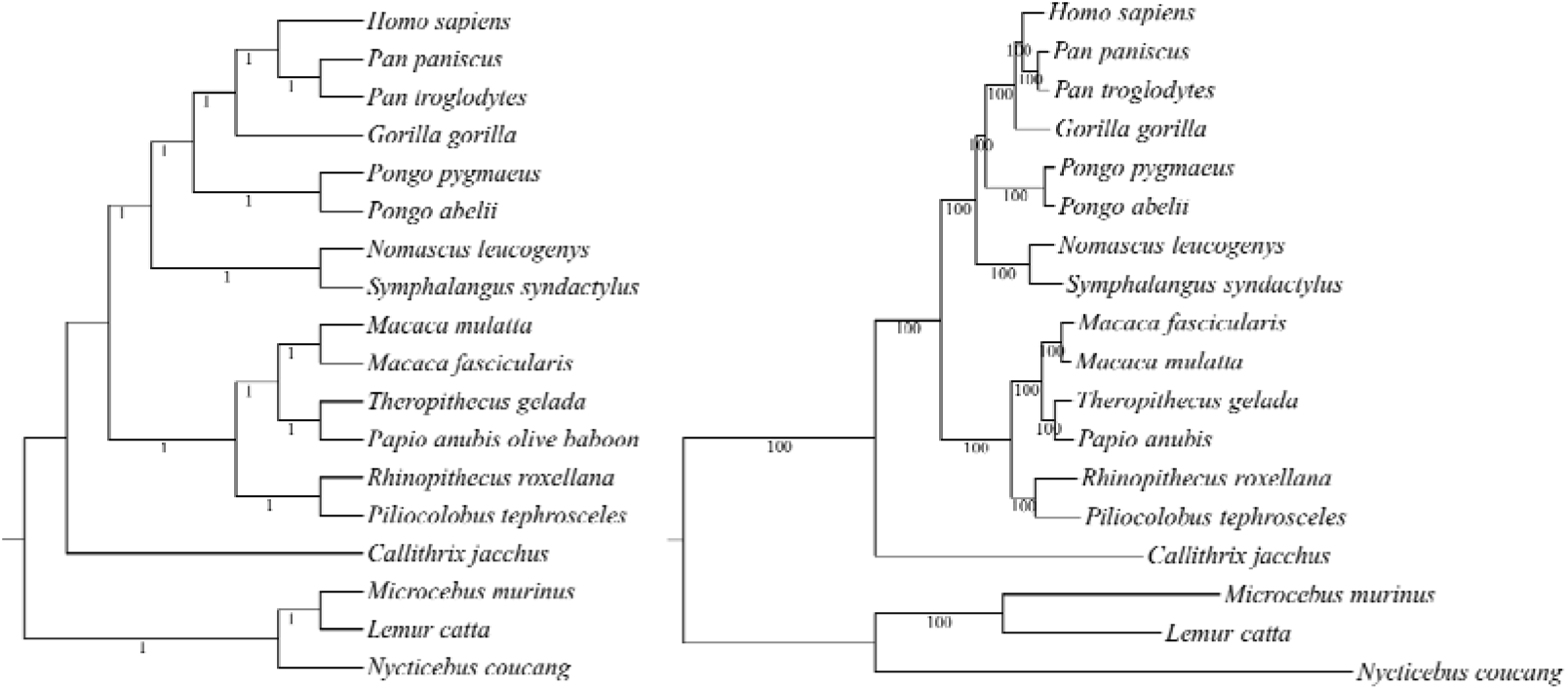
Coalescent-based (left) and concatenation-based (right) phylogenetic tree of 18 primate species.

**Supplemental figure 3.**
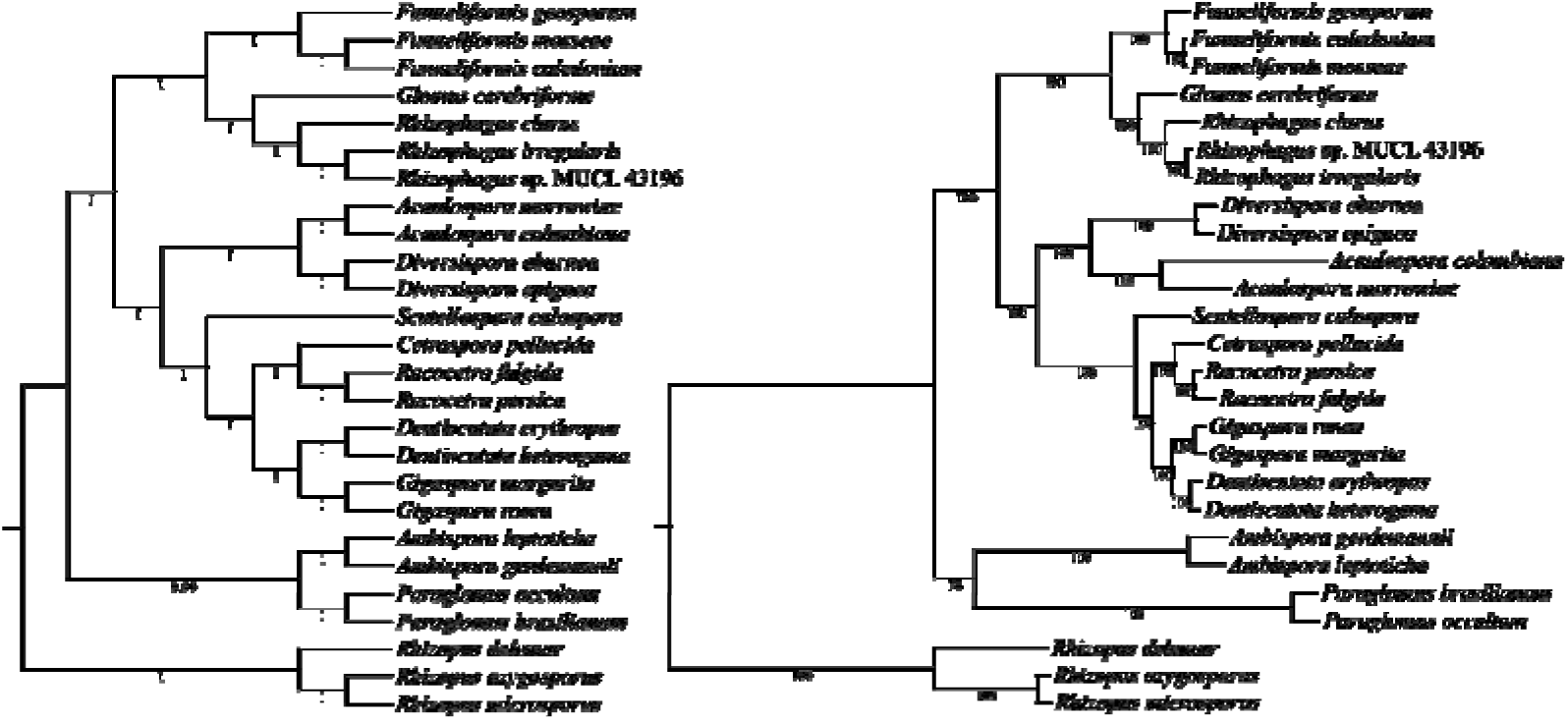
Coalescent-based (right) and concatenation-based (left) phylogenetic tree of arbuscular mycorrhizal fungi.

**Supplemental figure 4.**
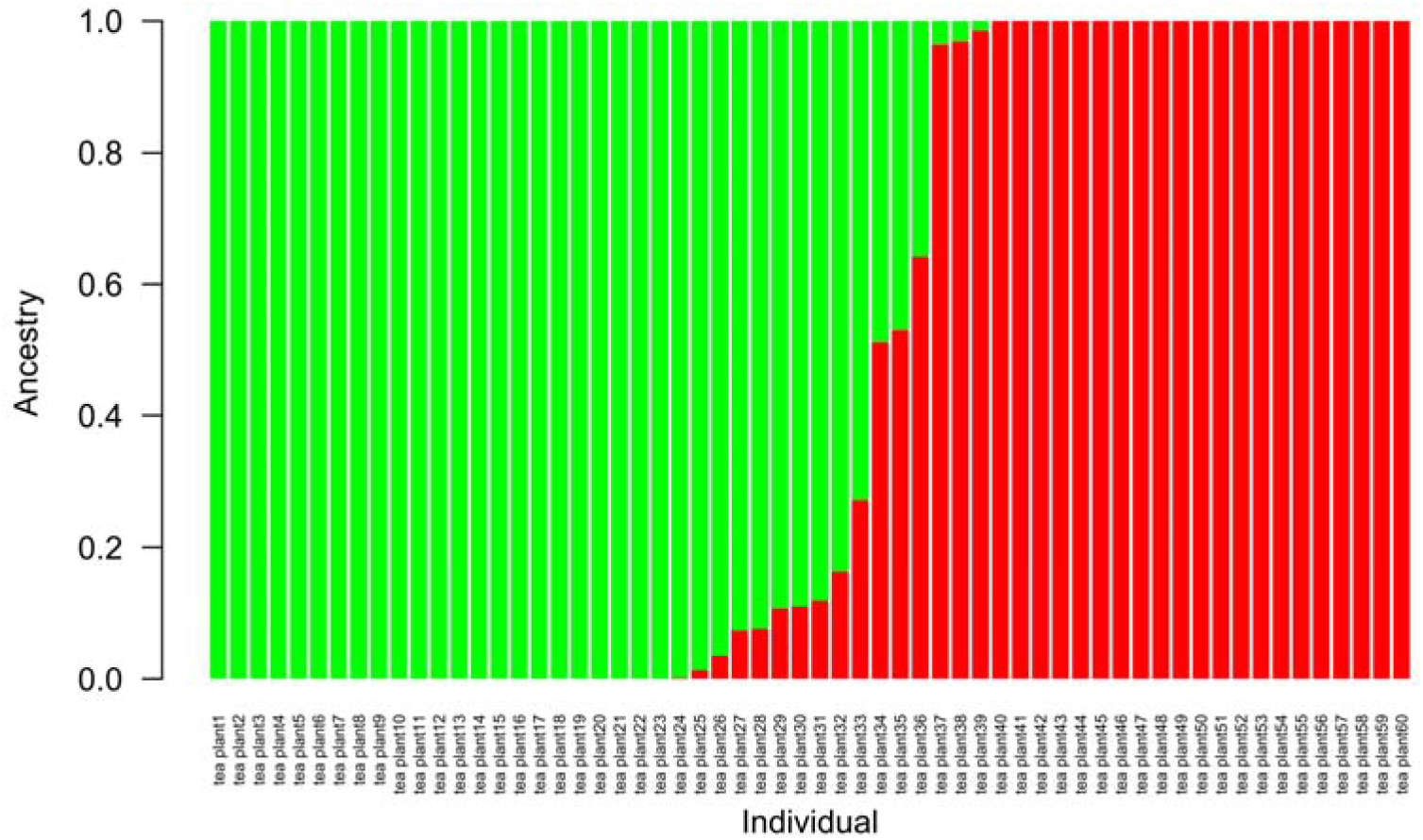
Population structure of 60 tea plants. CV error (K=2): 0.45470.

**Supplemental figure 5.**
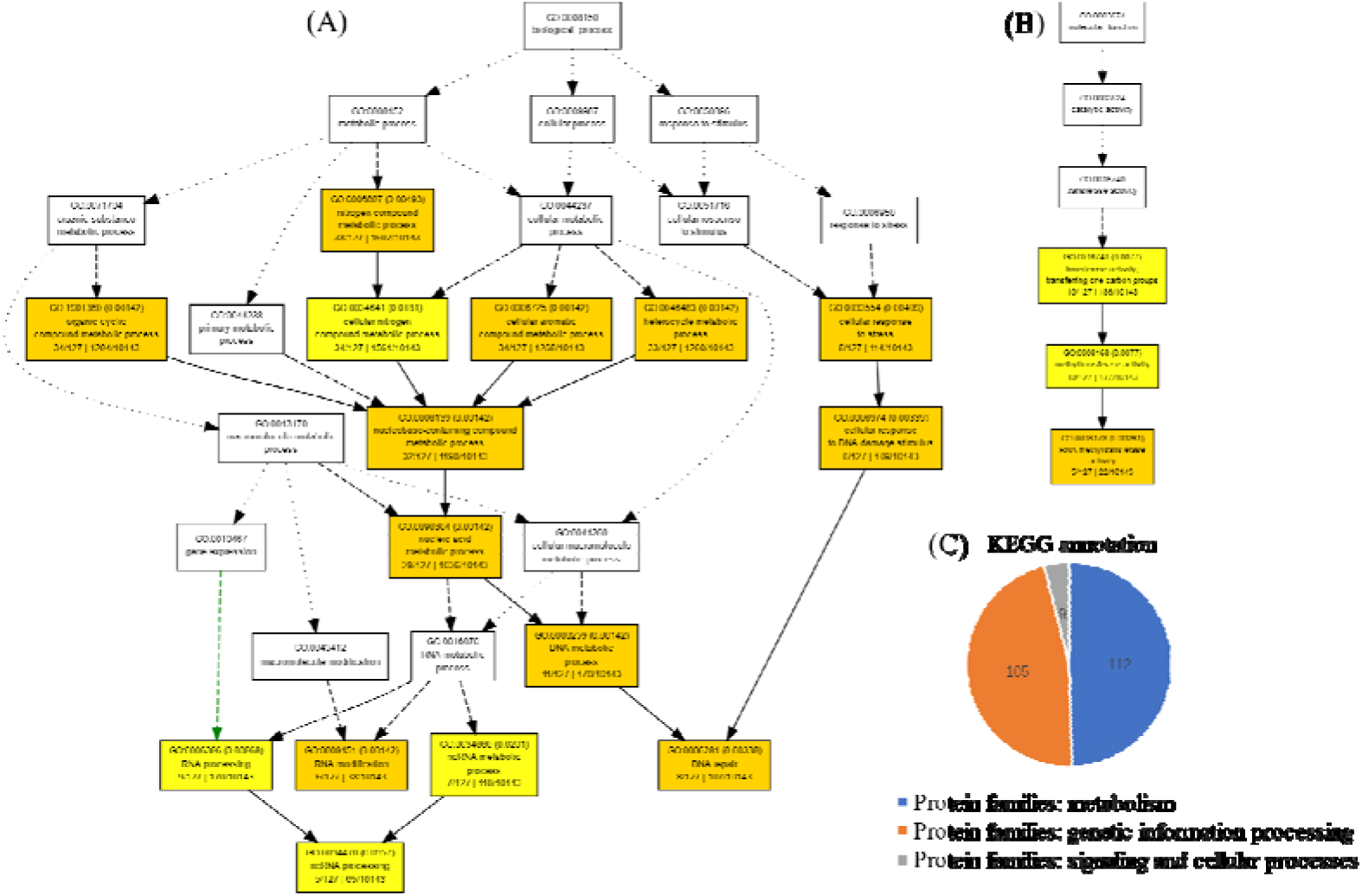
GO enrichment and KEGG annotation of low-copy BUSCO genes in Amborella trichopoda. (A) GO enrichment of BUSCO genes in biological process; (B) GO enrichment of BUSCO genes in molecular function; (C) KEGG annotation statistics of BUSCO genes.

**Supplemental Table 1:**
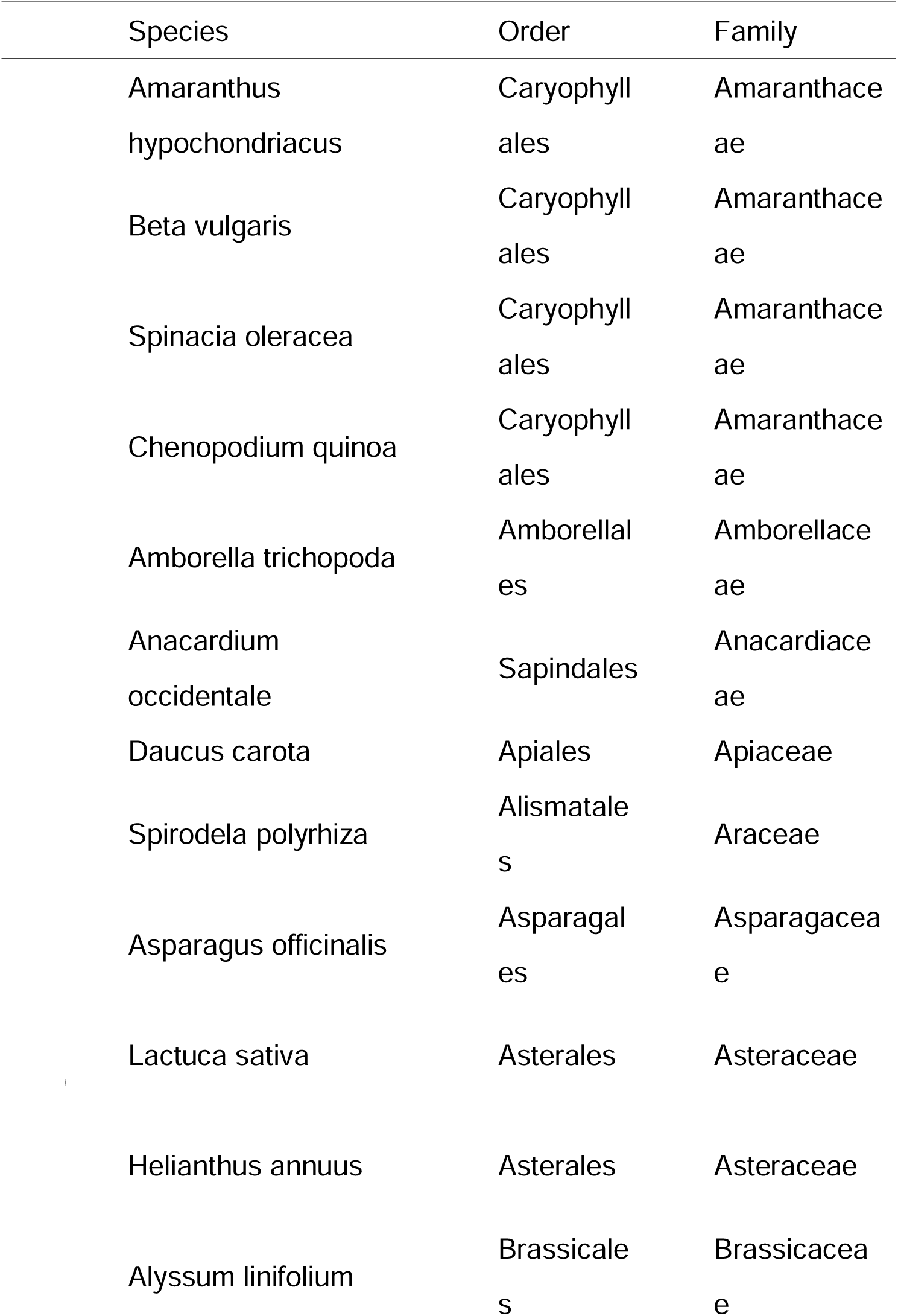
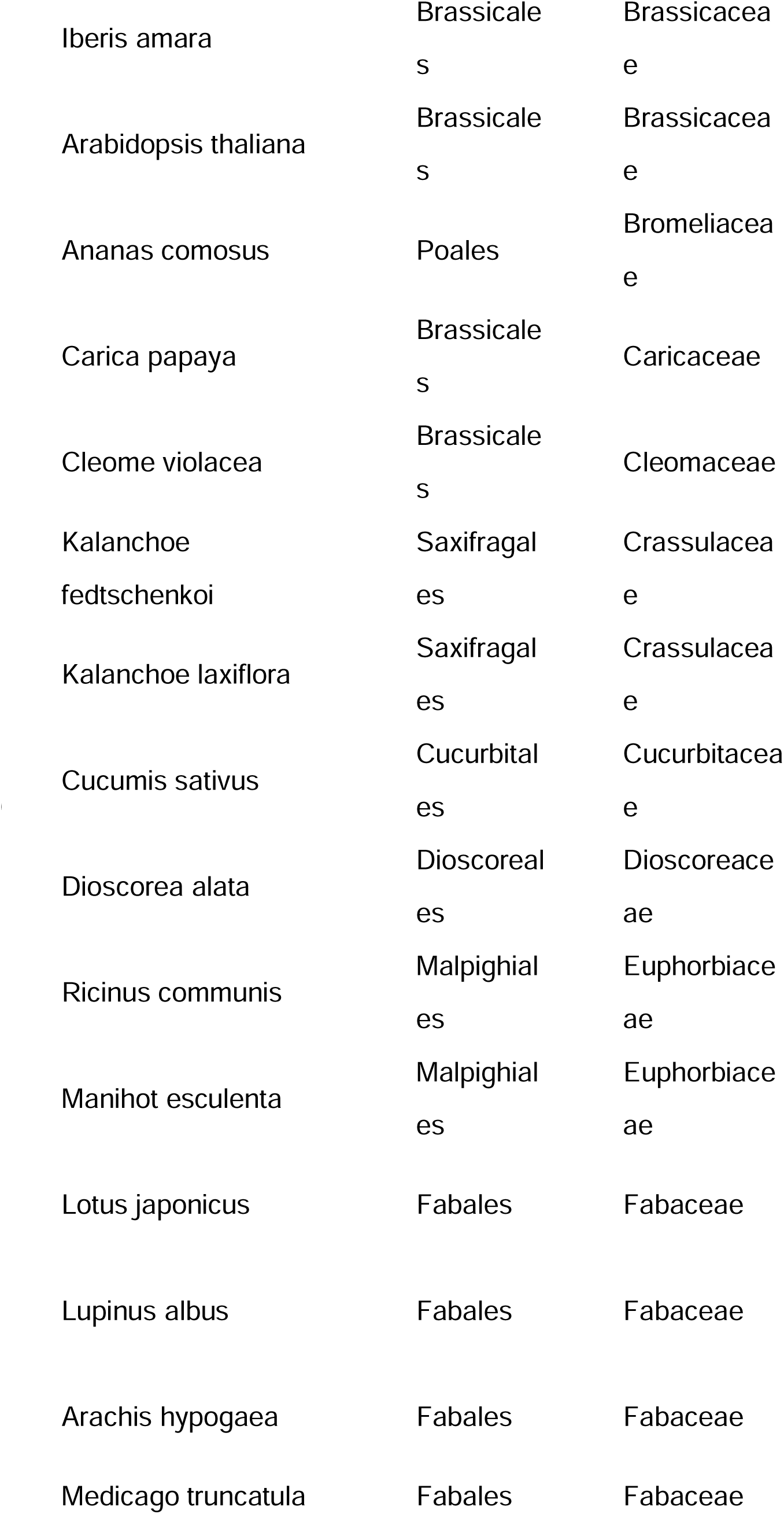
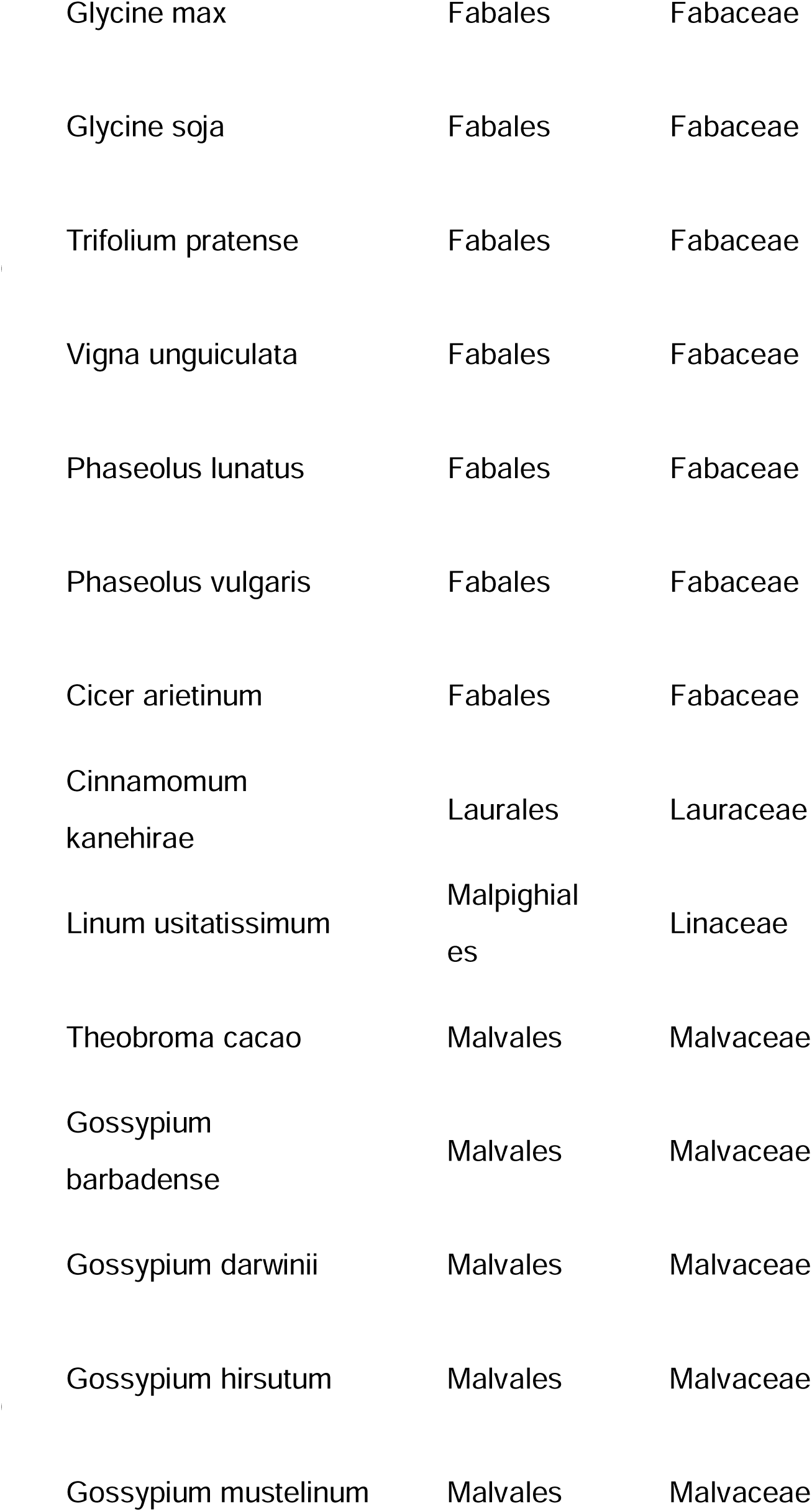
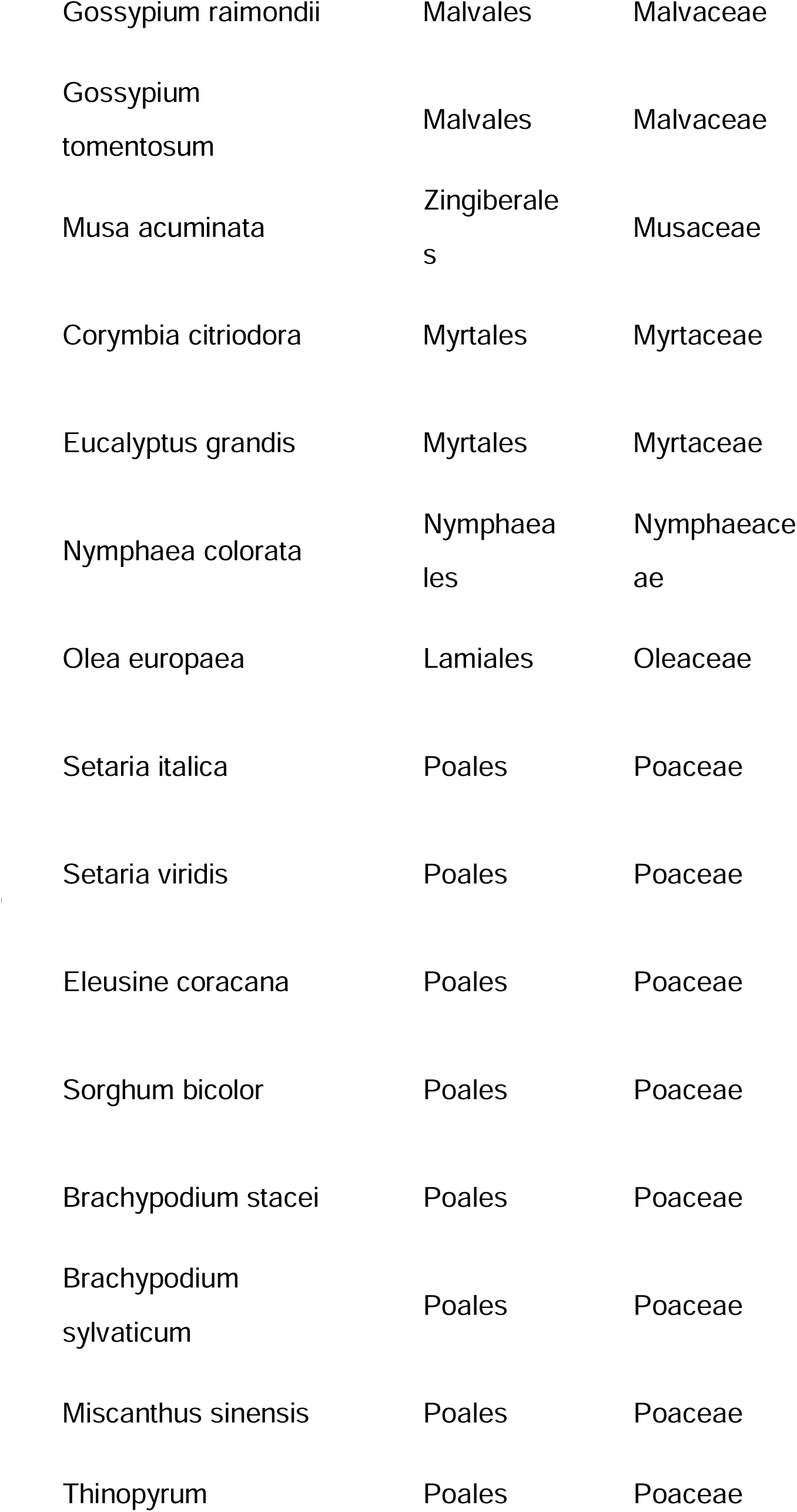
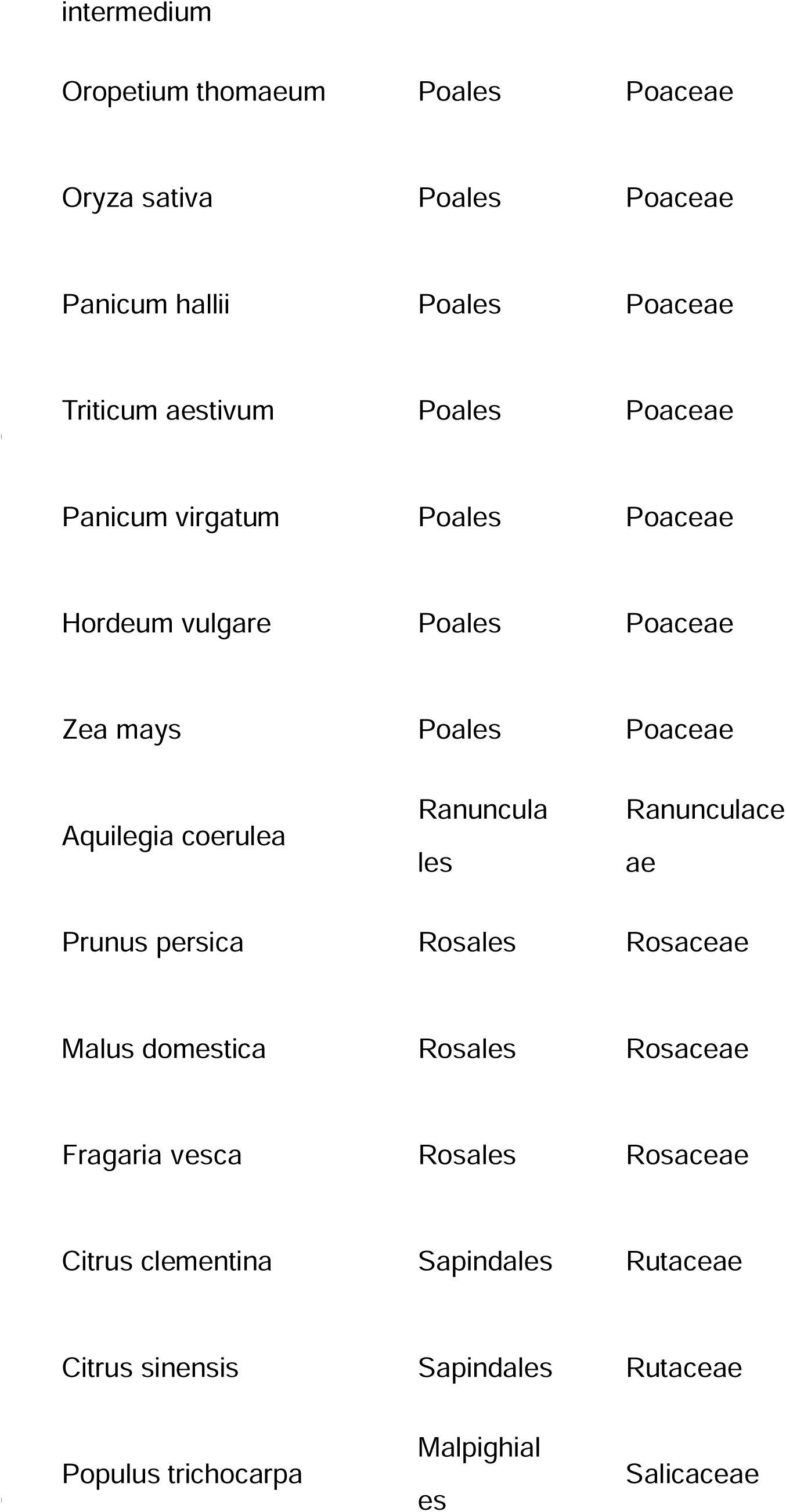
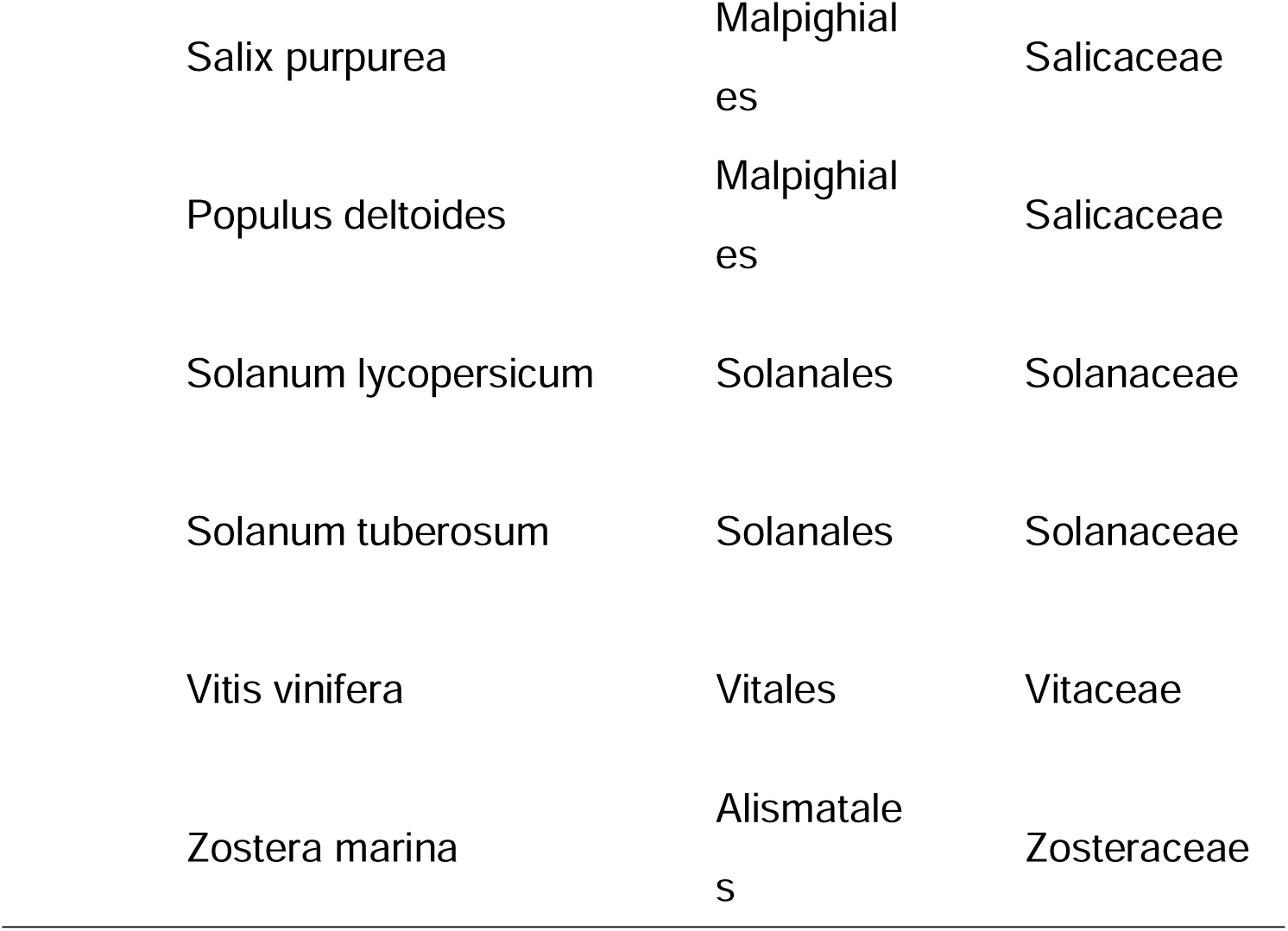
CDS data downloaded from Phytozome for Angiosperms.

**Supplemental table 2.**
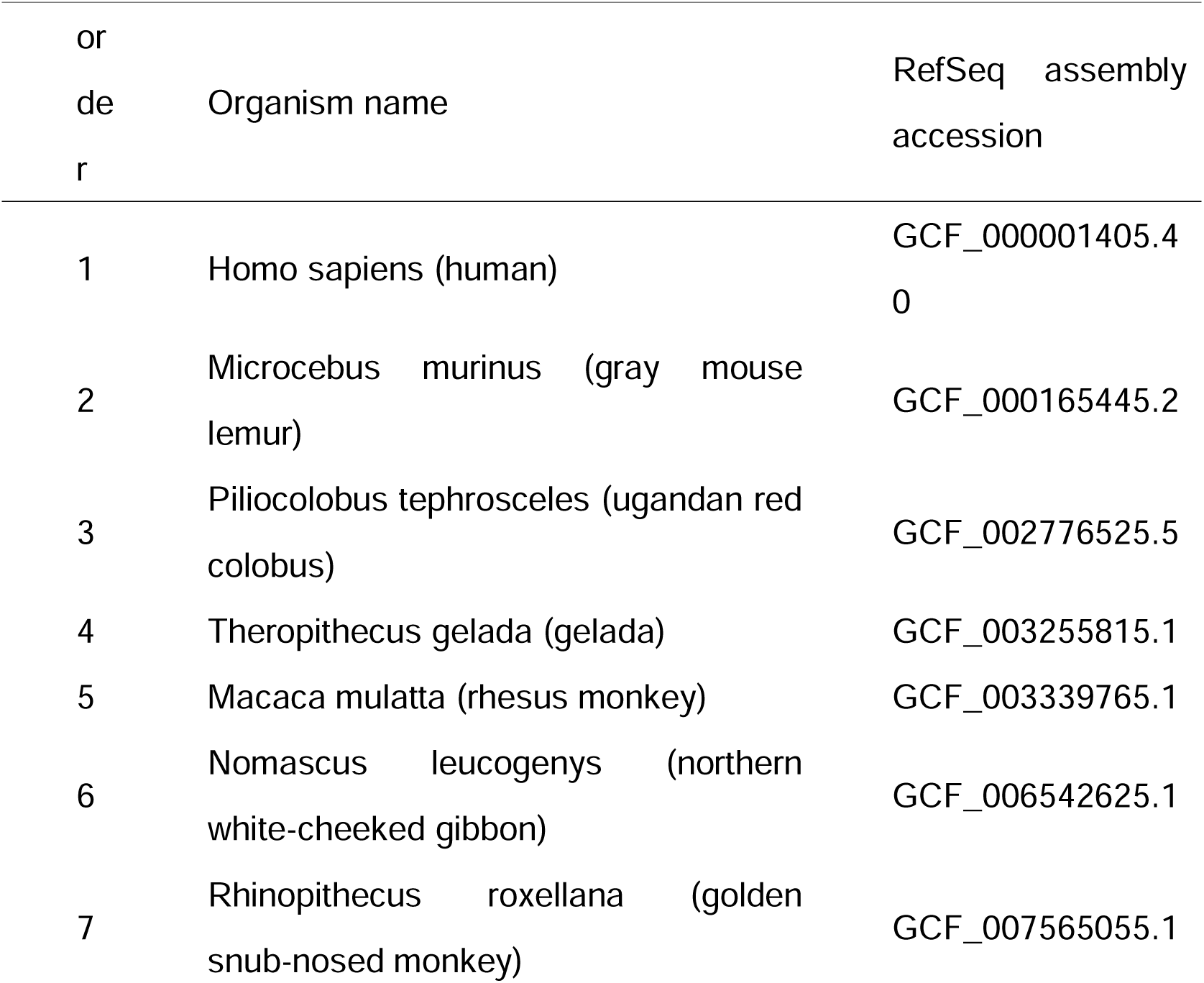
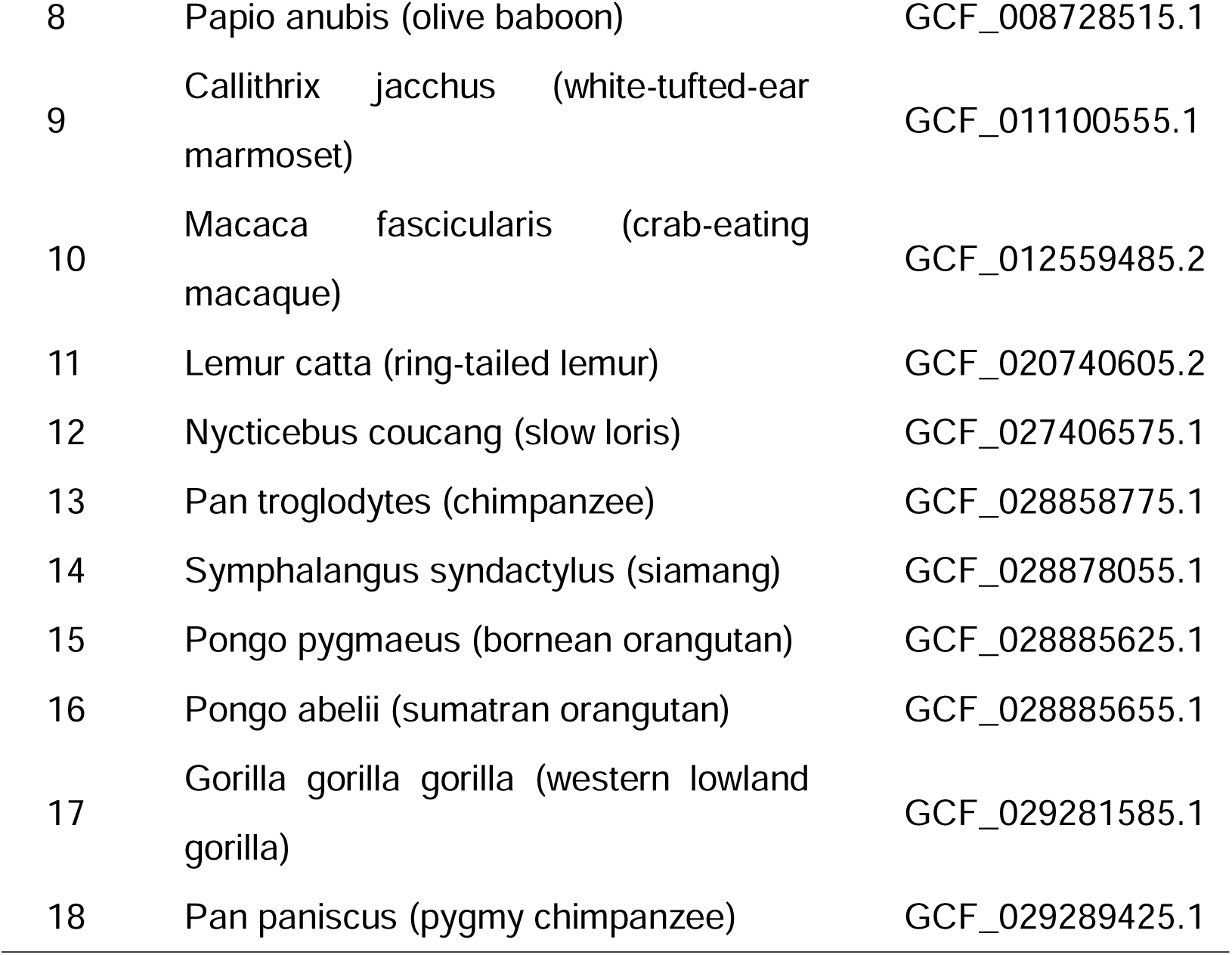
Details of CDS data downloaded for Primates.

**Supplemental table 3.** Details of CDS data downloaded for 23 species of arbuscular mycorrhizal fungi and three species of Mucoromycota.

| order | Organism name | GenBank accession |
| --- | --- | --- |
| 1 | Rhizopus delemar | GCA_000149305.1 |
| 2 | Rhizophagus irregularis | GCA_000439145.3 |
| 3 | Rhizopus microsporus | GCA_002708625.1 |
| 4 | Rhizopus azygosporus | GCA_003325435.1 |
| 5 | Diversispora epigaea | GCA_003547095.1 |
| 6 | Rhizophagus sp. MUCL 43196 | GCA_003549995.1 |
| 7 | Glomus cerebriforme | GCA_003550305.1 |
| 8 | Rhizophagus clarus | GCA_015698045.1 |
| 9 | Ambispora leptoticha | GCA_910591765.1 |
| 10 | Dentiscutata heterogama | GCA_910591775.1 |
| 11 | Funneliformis caledonium | GCA_910591825.1 |
| 12 | Acaulospora morrowiae | GCA_910591835.1 |
| 13 | Cetraspora pellucida | GCA_910591915.1 |
| 14 | Ambispora gerdemannii | GCA_910591945.1 |
| 15 | Dentiscutata erythropus | GCA_910591965.1 |
| 16 | Funneliformis mosseae | GCA_910592005.1 |
| 17 | Diversispora eburnea | GCA_910592015.1 |
| 18 | Gigaspora rosea | GCA_910592045.1 |
| 19 | Acaulospora colombiana | GCA_910592055.1 |
| 20 | Racocetra fulgida | GCA_910592135.1 |
| 21 | Paraglomus occultum | GCA_910592205.1 |
| 22 | Gigaspora margarita | GCA_910592245.1 |
| 23 | Racocetra persica | GCA_910592285.1 |
| 24 | Scutellospora calospora | GCA_910592325.1 |
| 25 | Paraglomus brasilianum | GCA_910592345.1 |
| 26 | Funneliformis geosporum | GCA_946474995.1 |

**Supplemental table 4.**
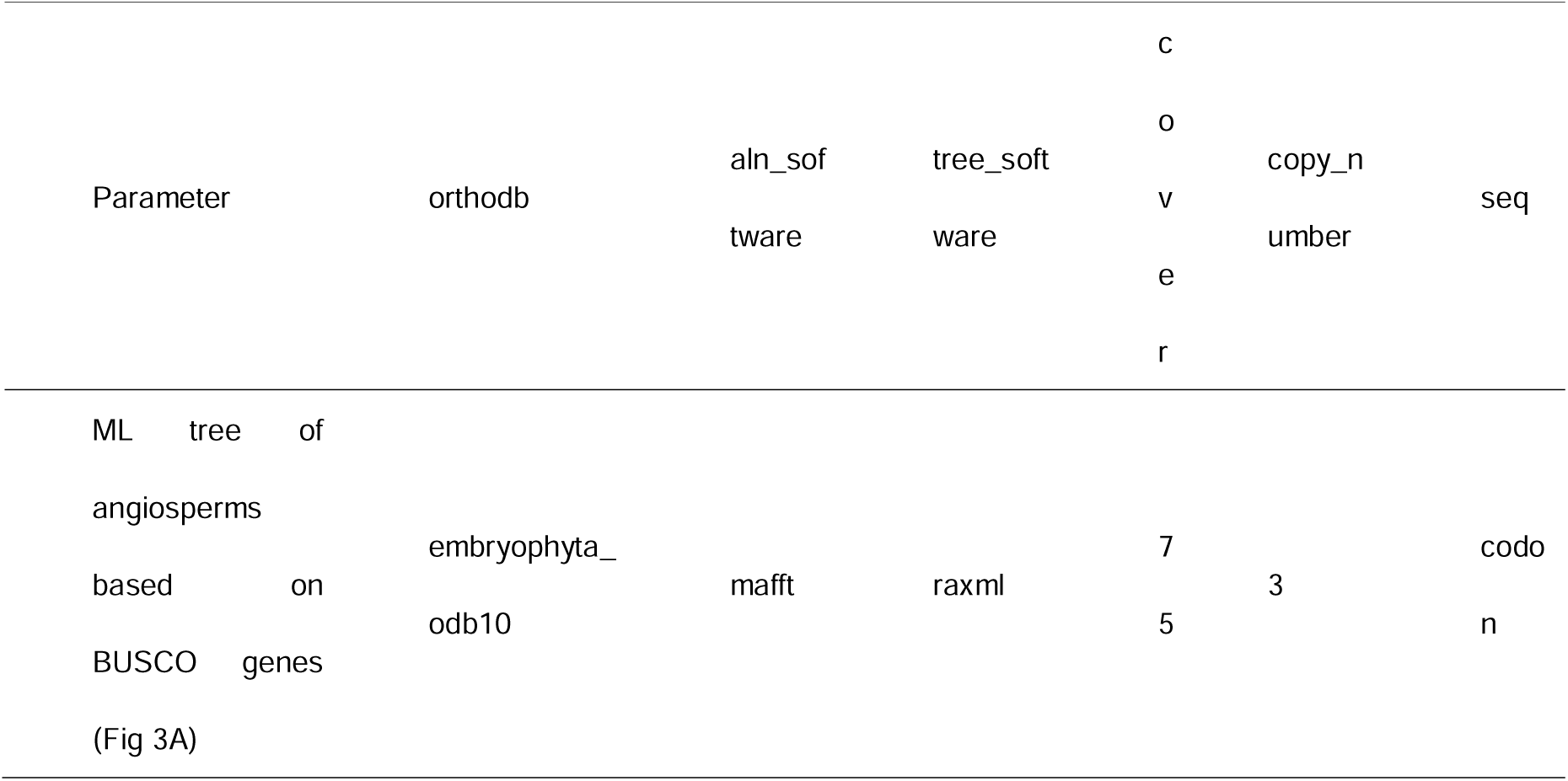
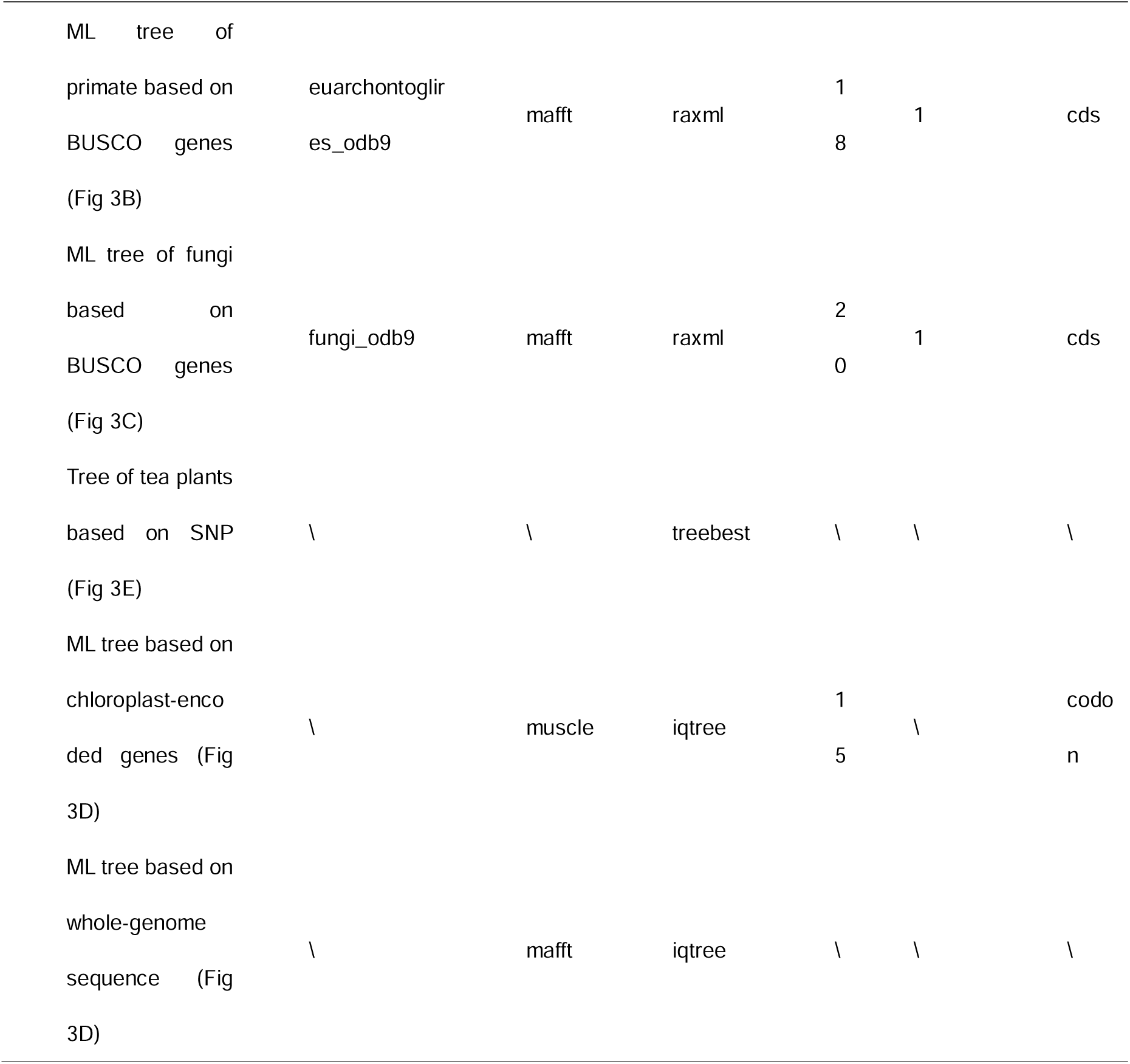
The detailed parameters of several tests in this paper.

## Notes

https://github.com/wangyayaya/PhyloForge

### Competing Interest Statement

The authors have declared no competing interest.

